# Fly motion vision is tuned to maximize signal energy transfer between mechanical input and sensor output

**DOI:** 10.1101/2024.03.29.587347

**Authors:** J. Sean Humbert, Holger G. Krapp, James D. Baeder, Camli Badrya, Inés L. Dawson, Jiaqi V. Huang, Andrew Hyslop, Yong Su Jung, Alix Leroy, Cosima Lutkus, Beth Mortimer, Indira Nagesh, Clément Ruah, Simon M. Walker, Yingjie Yang, Rafal W. Żbikowski, Graham K. Taylor

## Abstract

Insects achieve agile flight using a sensor-rich control architecture whose embodiment eliminates the need for complex computation. For example, their visual systems are tuned to detect the optic flow associated with specific self-motions, but what functional principle does this tuning embed and how does it facilitate motor control? Here we test the hypothesis that evolution co-tunes physics and physiology by aligning an insect’s sensors to its dynamically-significant modes of self-motion. Specifically, we show that the tuning of the blowfly motion vision system maximizes the flow of signal energy from gust disturbances and control inputs to sensor outputs, jointly optimizing observability and controllability. This evolutionary principle differs from the conventional engineering-design paradigm of optimizing state estimation, with implications for novel robotic systems combining high performance with low power-consumption.

## Main text

Like fifth-generation fighter aircraft and small multi-rotors, flies and other insects are inherently unstable in flight. This makes them highly maneuverable, but demands exquisite control. Technological and biological systems alike achieve this by combining information on motor input and sensor output (*1*) with an internal model of their dynamics (*2*), enabling them to observe and control their motion state in the face of disturbances. But whereas modern fly-by-wire control systems operate a computationally-intensive control architecture with recursive state estimation and a small number of sensors and actuators, insects have evolved a computationally-efficient control architecture with parallel processing and many sensors and actuators (*3*). For instance, the blowfly *Calliphora* fuses the output of ~10^5^ sensory cells to provide information on its self-motion, and uses at least 26 steering muscles to control its wingbeat. Yet, it weighs a mere 10^−4^ kg and consumes only 10^−2^ W of metabolic power in flight. The sensor-rich control architecture of insects may therefore point to a novel flight control paradigm in which specialised sensing avoids the need for generalised computation (*4*), but the underlying functional principle has yet to be identified (*5*).

One possibility is that an insect’s sensors are tuned to produce signals that directly correspond to excitation of the natural modes of motion characterising its flight dynamics (*3*). This principle, called the “mode-sensing hypothesis” (*3, 5*), might serve to reduce computational complexity through the use of embodied sensing. Under this hypothesis, an insect’s sensors need only detect the characteristic patterns of self-motion that are excited by gust disturbances and control inputs (*6*), rather than having to provide a calibrated measurement of some general physical quantity such as angular velocity or airspeed (*3*). The mode-sensing hypothesis accords with the broader observation that biological sensors are exquisitely sensitive to change but poor at measuring absolutes (*3*), even varying their gain according to the behavioral state of the animal (*7*). It might also explain why the descending neurons that relay sensory information downstream to the flight motor fuse information from multiple sensory modalities, because an insect’s flight dynamics are characterised by coupled rotational and translational motions that stimulate multiple sensory systems simultaneously (*3*).

Here we test whether the mode-sensing hypothesis explains the tuning of the fly motion vision system, which is currently the best-understood example of a deep neural network in nature (*8–10*). Visually-oriented animals including flies detect the wide-field optic flow stimuli experienced during self-motion by correlating local changes in luminance across neighboring photoreceptors before pooling this information globally. In flies, this operation is implemented by an array of elementary movement detectors whose responses are pooled by the lobula plate tangential cells (LPTCs) that form the output layer of the motion vision pathway. The LPTCs’ response characteristics are best known for *Calliphora* (*11*), functioning as matched filters (*12–14*) tuned to detect specific patterns of optic flow associated with particular combinations of rotational and translational selfmotion (*15, 16*). Any global tuning principle relating to the fly’s self-motion must therefore be embedded in the synaptic distribution and weighting of the dendritic inputs to its LPTCs, which are the unique level of the network at which information is pooled from across the optical array.

Each LPTC is tuned to detect some specific combination of rotation and translation defining a preferred direction of self-motion, which the mode-sensing hypothesis predicts should correspond to one or more dynamically-significant directions in the insect’s state space (*3*). In principle, this hypothesis can be tested by analyzing a suitable model of the insect’s flight dynamics (*5*), but no such model has yet been elaborated for *Calliphora*. Moreover, although rigid-body flight dynamics models (*17*) have been successfully developed for several other insect species (*18*), they do not usually attempt to model the output of the insect’s sensory system and do not accurately capture the detailed changes in wing kinematics involved in its flight control. Likewise, although recent neuromechanical models of insects are expressly designed to model the sensorimotor control of their behavior in a lifelike fashion (*19, 20*), few have yet attempted to model flight in a biomechanically accurate manner (*21*). Even then, as simulation models, these neuromechanical models are not designed to enable the abstraction of control-theoretic principles as is our aim here.

Our approach is therefore as follows: (i) to capture the dynamic mapping from mechanical input to sensor output analytically, which we achieve by creating a novel state-space model of blowfly flight dynamics and control; (ii) to identify the most dynamically-significant directions in the insect’s state space, which we accomplish by analyzing our state-space model using advanced control-theoretic tools called Gramians; and (iii) to test whether the preferred directions of the LPTCs correspond to these dynamically-significant directions more closely than expected by chance. Based on the strong correspondences that we identify, we conclude that blowfly motion vision is tuned to maximize the transfer of signal energy from control inputs and gust disturbances to sensor outputs via the system state (Fig. 1). This end-to-end tuning principle optimizes the observability of the system state jointly with its controllability and disturbance-sensitivity, which differs from the conventional engineering-design paradigm of placing sensors so as to optimize state estimation by maximizing observability alone (*22,23*). The evolutionary principle that we identify of tuning sensors to optimize observability jointly with controllability and disturbance sensitivity has important applications to the design of vehicles and robotic systems combining high performance with low computational load and low power consumption (*24*).

**Figure 1:**
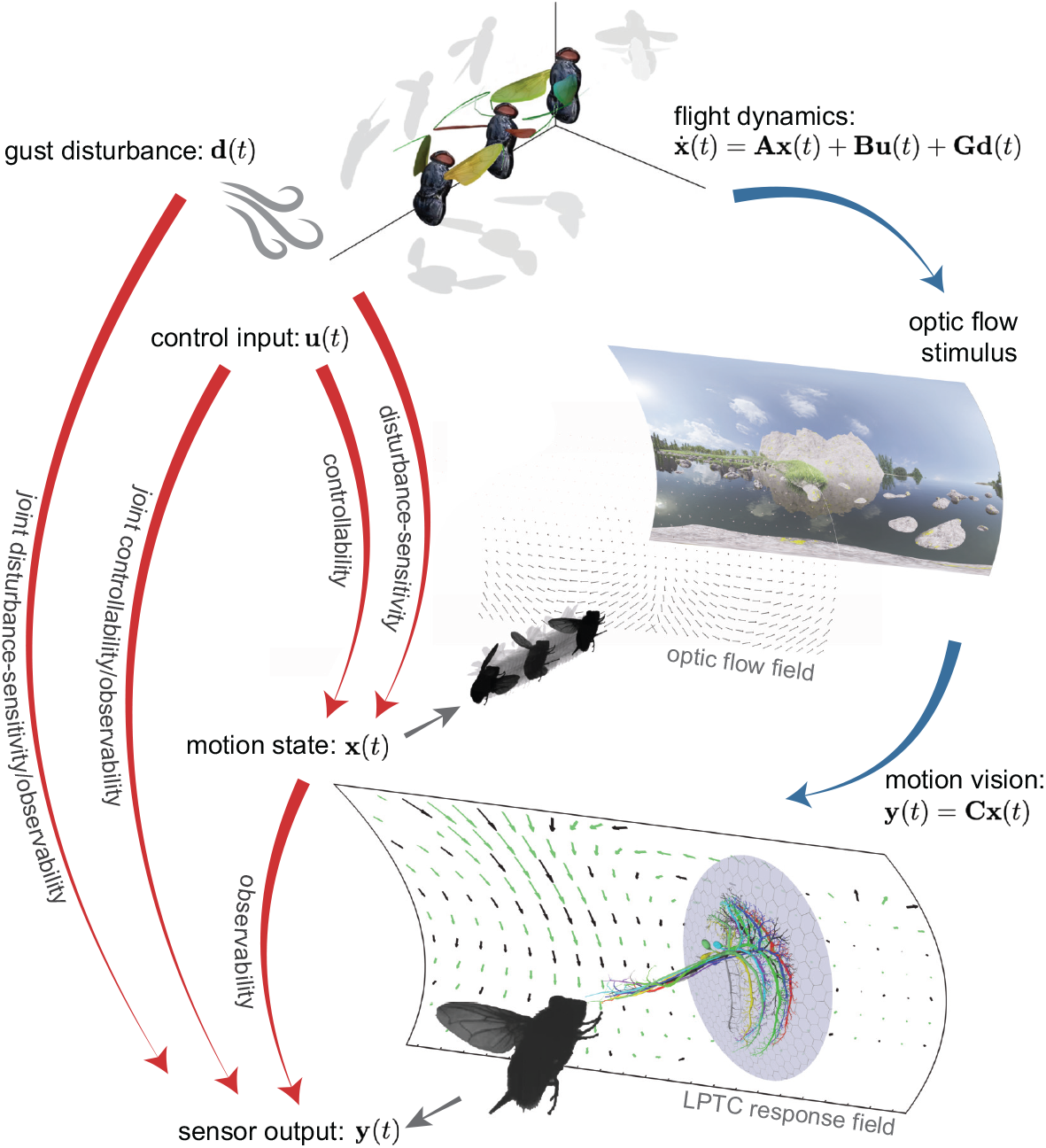
Signal energy transfer in insect flight. The mechanistic relationships illustrated on the right of the figure (blue arrows) are summarised by the signal energy flows shown on the left (red arrows). We characterise these mechanistic relationships by developing a state-space model of insect flight (Eqs. 1–2), and characterise the associated signal energy flows using the special matrix structures called Gramians that we derive from this model (Eqs. 3–4). Blue arrows: Control inputs **u**(*t*) and gust disturbances **d**(*t*) produce changes in the insect’s motion state **x**(*t*) described by the differential equations characterising its flight dynamics. This self-motion generates sensory stimuli including an optic flow field. The output layer of the blowfly motion vision system comprises a set of lobula plate tangential cells (LPTCs), each of which is matched to detect an optic flow field associated with a specific self-motion, yielding a sensor output **y**(*t*) related to the motion state **x**(*t*) of the fly. Red arrows: Signal energy from control inputs and gust disturbances is stored in the system state, so the system’s controllability and disturbance-sensitivity are maximized by maximizing signal energy storage. Stored signal energy is released through the evolution of the system state and retrieved at the sensor outputs, so the system’s observability is maximized by maximizing signal energy retrieval. A system that balances signal-energy storage/retrieval jointly maximizes observability and controllability or disturbance-sensitivity, and thereby maximizes the flow of signal energy from control inputs or gust disturbances to sensor outputs via the system state. Neuroanatomical image of LPTCs adapted from (*25*).

### Modeling approach

We begin by elaborating the novel flight dynamics model that we develop to characterize observability, controllability and disturbance-sensitivity in blowfly flight (Fig. 1). The simplest possible state-space model describing the dynamic mapping from mechanical input to sensor output linearizes an insect’s six degrees of freedom of rigid-body motion about some equilibrium flight condition (*17*) to yield the linear time-invariant equations: state equation:

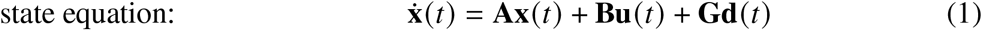

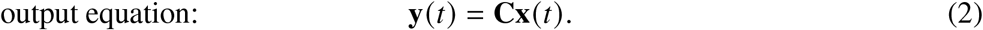

Here, all the forces and moments are assumed to be represented by their wingbeat averages, which is reasonable if the characteristic timescales of any unstable or oscillatory modes of motion are at least an order of magnitude longer than the wingbeat (*17*). This assumption is true of other flies (*18*), and we validate it directly here for *Calliphora* having first parameterized the model.

The state equation (Eq. 1) models the insect’s flight mechanics, and is parameterized by the system matrix **A** characterizing the insect’s natural response to perturbations in its motion state **x**. The control matrix **B** and disturbance matrix **G** characterize the insect’s forced response to control inputs **u** and aerodynamic disturbances **d**, respectively. The output equation (Eq. 2) models how the resulting self-motions map to the insect’s sensor output, and is parameterized by the output matrix **C** characterizing the physiological mapping from the insect’s motion state **x** to its sensor outputs **y**. Together, these two matrix equations describe the transfer of signal energy from control inputs and disturbances to sensor outputs via the system state (Fig. 1). Simplified versions of the state equation have been parameterized for a few other species (*18*), but these have not been coupled to an output equation modeling the resulting sensory output and they have not been founded on an accurate model of the kinematic inputs used in flight control. Moreover, to the best of our knowledge, there is no existing flight dynamics model for *Calliphora*. Given our model’s novelty and its centrality to the conclusions that follow, we therefore begin by detailing how we parameterize Eqs. 1–2 empirically in relation to a blowfly’s visual physiology and flight physics. We then analyze this model using advanced control-theoretic methods to identify the functional principle that the tuning of the blowfly’s motion vision system embodies.

### Visual physiology

We begin by characterizing the motion vision system whose tuning we aim to explain, providing the reader with a brief description of its anatomy and physiology, before using new and published electrophysiological recordings to parameterize the output equation (Eq. 2).

#### Visual output vector

The output layer of the fly motion vision system is formed by a set of wide-field optic-flow sensitive neurons called the lobula plate tangential cells (LPTCs). A subset of the LPTCs in *Calliphora* is tuned to respond specifically to self-motion stimuli, including the ten vertical system (VS) cells {VS1–VS10} and three horizontal system (HS) cells {HSN, HSS, HSE} of each optic lobe (*11, 15, 26, 27*). The VS- and HS-cells arborize ipsilaterally, yet some of their response fields extend across both visual hemispheres (Fig. 2B), which is important to distinguishing rotational from translational self-motion (*3*). This binocularity is made possible by a complex coupling arrangement (Fig. S5) in which heterolateral LPTCs called V-cells {V1, V2, Vx} and H-cells {H1, H2, Hx} relay output from the contralateral optic lobe (*11, 27, 28*). The binocular VS- and HS-cell outputs are ultimately combined with output from other sensory modalities involved in flight control by descending neurons that relay sensory information to the wing, leg, and neck motor systems (*29, 30*). Hence, whereas the VS- and HS-cells form the output layer of each optic lobe, the heterolateral V- and H-cells form a shallow hidden layer that is critical to the function of this bilaterally symmetric deep neural network (Fig. S5). To allow us to analyse their respective functions, we use the characteristic responses of all 19 cells in each of the two mirror-symmetric optic lobes to form the 38 elements of the output vector **y**. To enable us to model this output vector **y** we must quantify the LPTC responses using a combination of new and existing electrophysiological recordings, as described in the next section.

**Figure 2:**
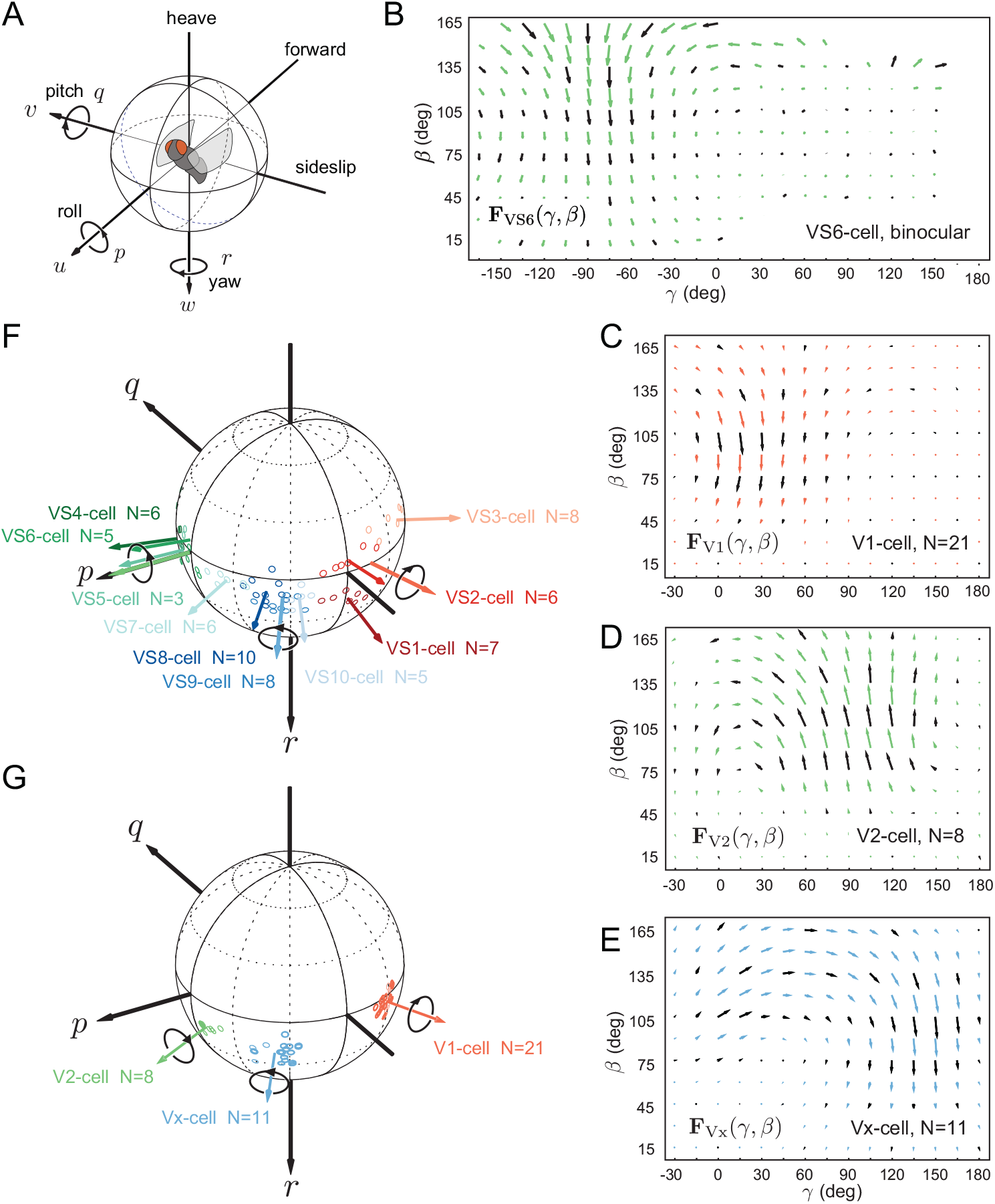
The fly motion vision system. **(A)** A flying insect has six degrees of freedom in rotation {*p, q, r*} and translation {*u, v, w*}. Its compound eyes sense self-motion using a deep network comprising an array of repeated elementary motion detectors whose outputs are pooled by the lobula plate tangential cells (LPTCs), comprising an output layer of 13 pairs of VS- and HS-cells connected by a hidden layer of at least 6 pairs of V- and H-cells coupling the left and right optic lobes. **(B)** Binocular response field of the left VS6-cell as a function of azimuth (*γ*) and elevation (*β*) in retinal coordinates. This closely resembles the optic flow associated with left-handed roll motion (*28*), and as the LPTCs only arborize ipsilaterally, the weak contralateral response visible here must be due to coupling by the heterolateral V-cells. **(C**,**D**,**E)** Newly-measured response fields of the right V-cells, where N denotes the number of individuals from which the recordings were pooled. **(F**,**G)** Preferred rotation axes of the left VS-(*15, 26*) and V-cells (new data); cells of the right optic lobe have responses that are mirror-symmetric to those of the left optic lobe.

#### Electrophysiological measurements

Characterization of the LPTCs’ electrophysiological responses to local image motion (Fig. 2B-E) reveals that each cell’s response field (i.e., the vector field describing its local motion sensitivity and local preferred directions) resembles a coherent optic flow stimulus associated with some specific combination of rotational and translational selfmotion (*16*). For example, it is well known that the VS-cells signal combinations of heave translation and roll or pitch rotation (*15, 26*) (Fig. 2F), whereas the HS-cells signal combinations of yaw rotation and sideslip or forward translation (*27*). The responses of the heterolateral LPTCs have been less well studied, so to complete our model of the blowfly motion vision system, we characterised the V1, V2, and Vx response fields of male and female flies experimentally (Fig. 2C-E). We did so by making extracellular recordings of the cells’ activity in response to local image motion, which we used to determine the spatial distribution of their local preferred directions and local motion sensitivity across the visual field (see Methods). These new data from both sexes complement and complete existing recordings made extracellularly from the V- and H-cells (*27, 31, 32*) and intracellularly from the VS, HS and Hx cells of females only (*15, 26–28*). Our results show no functional differences between males and females, and demonstrate that the preferred rotation axes of V1, V2 and Vx (Fig. 2G) each coincide with different subsets of the VS-cells (Fig. 2F; see also Fig. S5). The correspondence between the response fields of the LPTCs and the optic flow fields associated with specific combinations of rotational and translational self-motion is striking, but the dynamical significance of these patterns has only been examined qualitatively to date (*3*).

#### Physiological modeling of the output equation

The response fields {**F**_*i*_} of the *i* = 1, …, 38 LPTCs of the left and right optic lobes are defined in retinal azimuth and elevation coordinates {*γ, β*} whose equatorial plane *β* = 0 is assumed to be held horizontal at equilibrium. The retinal coordinate system is assumed to rotate with the body such that the ray defining its origin coincides with the *x*-axis used in our flight dynamics modeling (Fig. 3). This approximation is valid for the small perturbations that we model, and is reasonable to the extent that compensatory head movements are driven optokinetically (*33, 34*) and therefore lag the body’s motion (*35*). The magnitude of the optic flow experienced during translational self-motion varies inversely with distance to the visual environment. To determine how the LPTCs are expected to respond to rigid-body motion, we assume that the insect is flying at the centre of a 2 m cube, although we relax this assumption later. To parameterize the output equation (Eq. 2) in this environment, we compute the partial derivative of the optic flow field 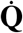 with respect to each element of the state vector **x** (see Methods). We then take the inner products of these matrices with the LPTC response fields {**F**_*i*_}, and use these to form the elements of the unilateral output matrix **C**′, whose 38 normalized rows describe the preferred directions of self-motion of the 19 mirror-symmetric pairs of LPTCs, treating the outputs of the left and right optic lobes separately (Table S7). Finally, because symmetric and asymmetric motions can be treated separately in our linearized flight dynamics model (see Fig. 3F), we restructure **C**′ to form a bilateral output matrix **C** whose 38 normalized rows represent the summed and differenced responses of the 19 mirror-symmetric pairs of LPTCs (see Fig. 4A below). This pairwise approach to combining the output of the left and right optic lobes is intended to represent the fact that each LPTC pair carries separable information on symmetric versus asymmetric motions, and makes no explicit assumptions on the actual downstream connections of the LPTCs, which is necessary because our knowledge of how the descending neurons combine this information remains incomplete (*36–38*).

**Figure 3:**
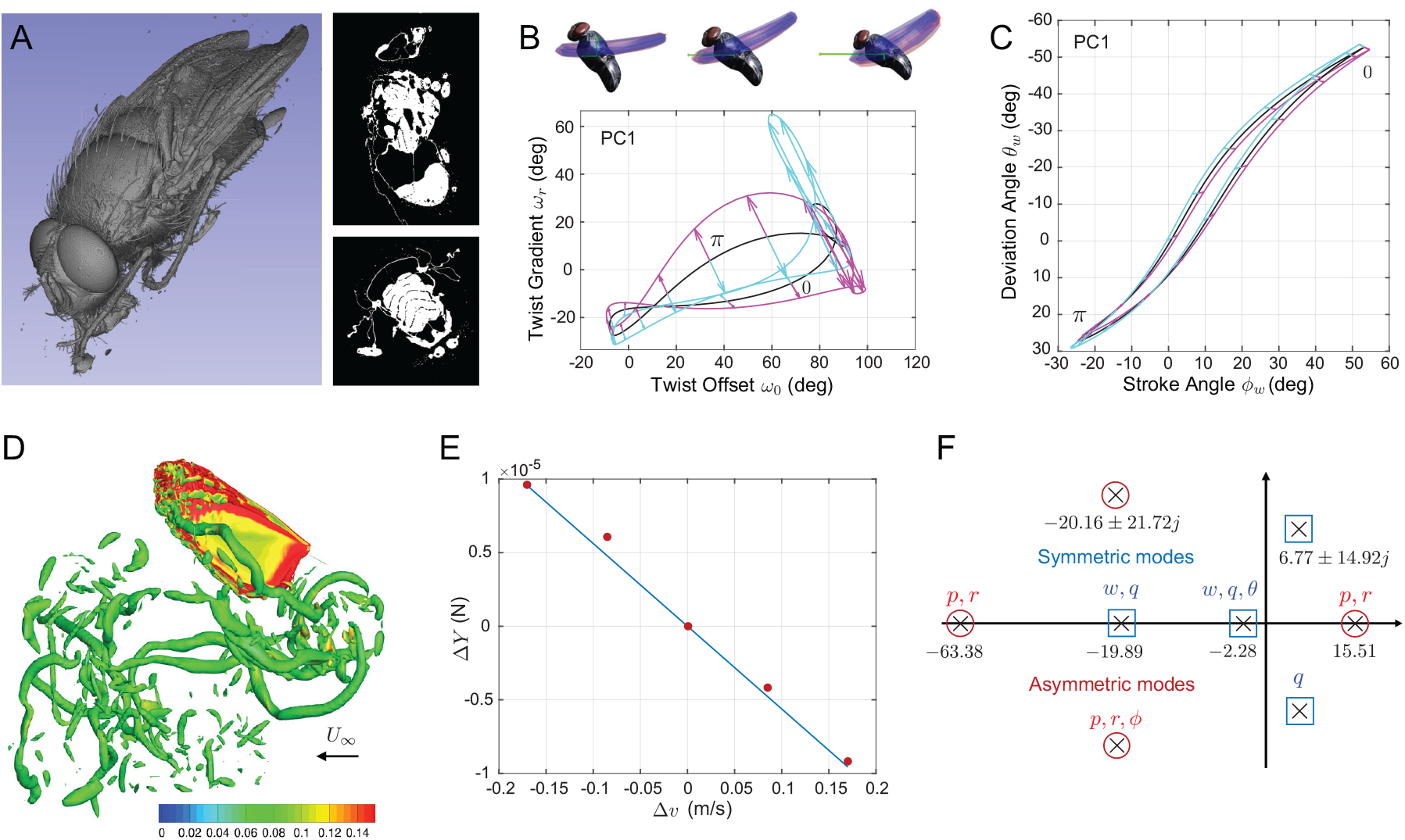
Modeling of blowfly flight physics. **(A)** We estimated the fly’s inertia tensor using synchrotron-based X-ray microtomography: images show a 3D rendering and longitudinal/transverse slices of a male blowfly. **(B**,**C)** We used high-speed videography to record the wing kinematics of free-flying blowflies over a range of flight speeds, and used functional principal components analysis to define a set of control inputs summarising the coupled variation in wing-twist (B) and tip (C) kinematics: phase portraits illustrate the reference wing kinematics (black) ±1 S.D. (cyan/magenta) in the first principal component (PC1). Note that PC1 involves coupled changes in stroke plane, stroke amplitude, and wing twist, which we may conceptualize as characterizing the result of the fly’s limit-cycle control of its wingbeat oscillation (*39*). **(D)** We used a Reynolds-averaged Navier Stokes solver to model how the aerodynamics vary with the kinematics; this image of the wing and wake shows vortex shedding at the end of the downstroke, visualized using iso-*Q* criterion surfaces (*Q* = 0.001) colored by vorticity. **(E)** We estimated the fly’s stability and control derivatives by regressing the wingbeat-averaged aerodynamic forces and moments on the perturbed states or control inputs for a single wing; this plot shows the zero-intercept regression of the change in lateral force with respect to the change in lateral velocity; control derivatives were estimated similarly by modeling the changes in the aerodynamic forces with respect to each modeled control input. **(F)** Eigenstructure of the system matrix: the eigenvalues of **A** are plotted in the complex plane, with labels denoting the dominant components of self-motion for the associated eigenvectors; note that there is a pair of unstable modes with positive real parts, indicating that the fly’s flight dynamics are inherently unstable.

**Figure 4:**
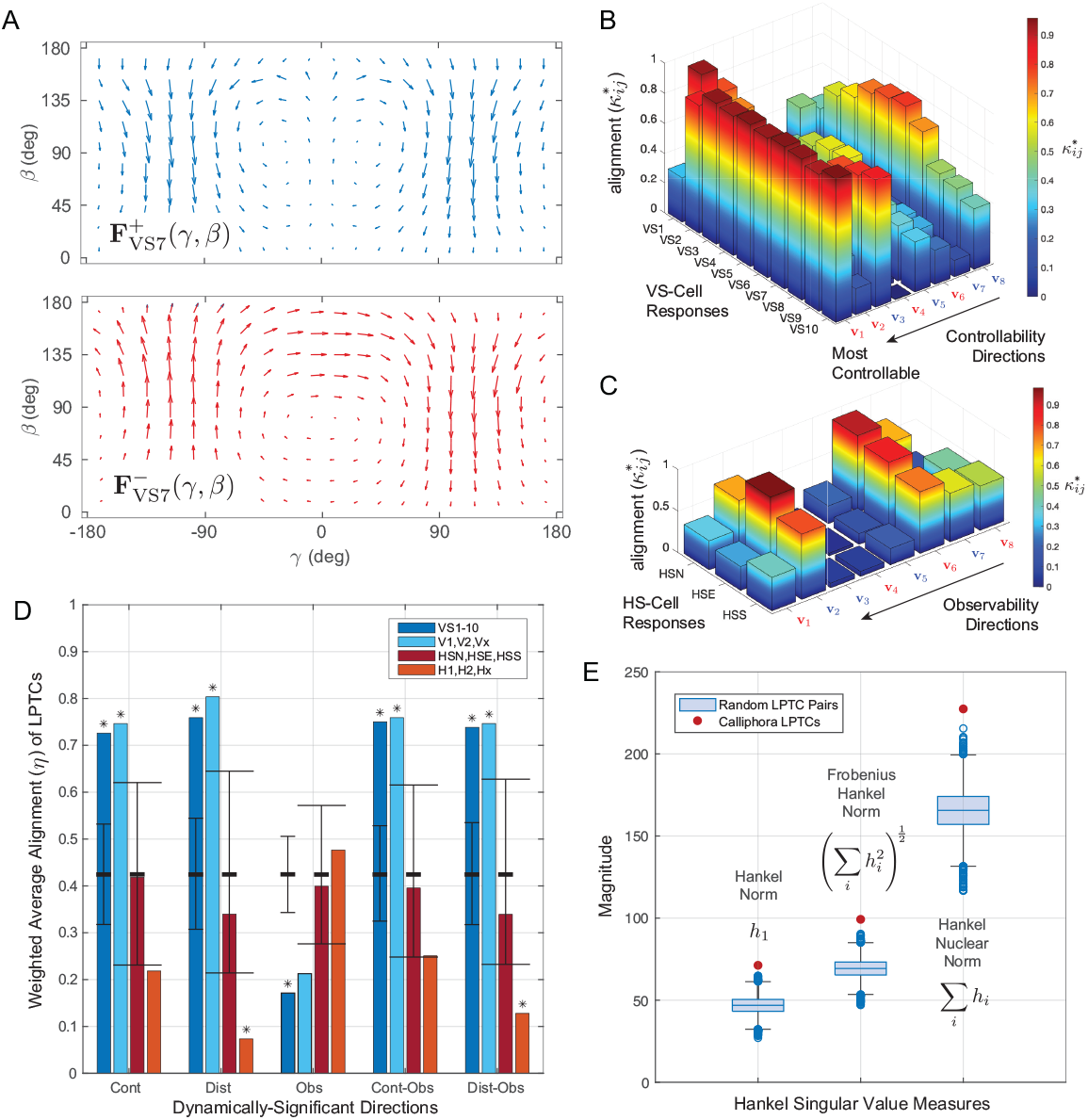
Functional principles of blowfly sensorimotor design. **(A)** Each LPTC pair carries separable information on symmetric versus asymmetric motion, which we illustrate by taking the sum (blue) or difference (red) of their response fields, shown here for VS7. **(B**,**C)** These response fields encode the cells’ preferred directions of rotational and translational self-motion, which the mode-sensing hypothesis predicts will be aligned to dynamically-significant directions in the insect’s state space. To test this, we quantified their alignment to the ordered symmetric (blue) or asymmetric (red) eigenvectors of the controllability and observability Gramians {**v**_1_, …, **v**_8_}, ranked by their dynamical significance. Collectively, the VS-cells strongly encode all of the most-controllable directions of symmetric/asymmetric motion, whereas the HS-cells strongly encode only the most-observable direction of symmetric motion. **(D)** Averaging over each LPTC sub-population, the V- and VS-cells strongly encode the directions that maximize controllability or disturbance-sensitivity both independently and jointly with observability; stars (*) denote statistical significance controlling the overall Type I error at *α* = 0.05. Note that no LPTC sub-population is aligned to the most-observable directions of motion, as would be conventional in engineering design. **(E)** To assess overall system performance, we computed three measures (the Hankel norm, the Frobenius-Hankel norm, and the Hankel Nuclear norm) of the Hankel singular values {*h*_1_, …, *h*_8_} characterizing signal energy flow through the system for all 19 LPTC pairs (red points) and compared these to the Hankel singular values of 100,000 randomly-generated sets of preferred directions (blue box plot; whiskers extend ± 2.7 S.D.). The blowfly’s Hankel singular values are far higher than expected by chance, demonstrating that the LPTCs’ tuning maximizes signal energy flow through the system. The first Hankel singular value provides an upper bound on signal energy flow from input to output, and is higher for the blowfly than for any of the randomly-generated systems.

### Flight physics

The parameterized output equation (Eq. 2) models how the insect’s six degrees of freedom of rigid-body motion motion are represented in the 38-dimensional output of its motion vision system. To understand how this sensor output responds to control inputs and aerodynamic disturbances, we must also model the insect’s flight dynamics by parameterizing the state equation (Eq. 1). Previous state-space models of insect flight control (*18*) have not attempted to identify the detailed control inputs that are available to the insect, so building a biologically-meaningful model requires the development of new analytical modeling approaches, as described in the sections below.

#### State and disturbance vector

The insect’s rigid-body flight dynamics (Fig. 2A) are described by the Newton-Euler equations of motion relating its linear and angular acceleration to the gravitational and aerodynamic forces and moments (*17*). For convenience, these vector quantities are defined in body-fixed stability axes whose *x*-axis is aligned to the flight velocity vector at equilibrium and whose *y*-axis is normal to the insect’s symmetry plane. It follows that the insect’s state vector **x** must contain complete information on its linear velocity **v** = [*u v w*]^*T*^ and angular velocity *ω* = [*p q r*]^*T*^ with respect to an inertial reference frame, together with information on the roll and pitch angles of the body {*ϕ, θ*}, which is needed to keep track of gravity as the insect rotates. For the linearized system in Eqs 1–2, these 8 elements of the state vector **x** are treated as small perturbations (*δ*) from level symmetric rectilinear flight at equilibrium, and are separated into their symmetric and asymmetric parts **x**_long_ = [*δu δw δq δθ*]^*T*^ and **x**_lat_ = [*δv δ p δϕ δr*]^*T*^, describing longitudinal and lateral motions, respectively. Bulk motion of the surrounding air mass produces the same relative airflow as translational or rotational self-motion, so we use analogous perturbation quantities to form the aerodynamic disturbances in the disturbance vector **d**.

#### Reference wing kinematics

A blowfly’s wingbeat is a complex three-dimensional limit cycle motion involving substantial aeroelastic deformation, driven by muscular forces applied at the wing root through one of the most complex linkages in the natural world (*40*). To capture this complexity, we used four high-speed video cameras to record the deforming wing kinematics of N=28 individuals over 274 flights at 3,800 fps, and used a voxel carving method (*41*) to identify the three-dimensional outline of the wings and estimate the pose of the body (Fig. 3B); see Methods. We measured the angular position of the wing tips in a body-fixed coordinate system (Fig. 3C), and estimated the torsional deformation of the wings under a linear twist distribution (Fig. 3B). We then fitted a Fourier series with linear trend to each of the 2,708 wingbeats that we recorded for either wing. We defined a set of reference wing kinematics for our aerodynamic modeling by averaging the Fourier coefficients over three wingbeats representing near-equilibriun flight. These three wingbeats were centered on the wingbeat that most nearly achieved level flight from within the subset of wingbeats associated with near-zero body acceleration (< 0.5 m s^-2^). For generality, we set the reference wingbeat frequency (*f*_*e*_ = 166 Hz), reference flight speed (*u*_*e*_ = 0.85 m s^-1^), and reference body pitch angle (*θ*_*e*_ = 30°) to their mean values over all of the wingbeats we had measured, and used these to model a reference condition of level forward flight.

#### Control input vector

Whereas the control inputs of an aircraft are known by design, and include simple mechanical quantities such as throttle settings and control surface deflection angles, insect wings are passive structures that lack discrete control surfaces. This makes it challenging to identify what inputs an insect’s control vector **u** should contain, but defining this in a biologically meaningful way is essential to any meaningful analysis of controllability. We therefore used functional principal components analysis (*42*) to summarize the empirical variation in the Fourier coefficients describing each of the 2,708 wingbeats that we had measured. This analysis decomposed the observed aeroelastic variation in the wing kinematics into a set of principal components (PCs) characterizing the dominant kinematic couplings (Fig. 3B-C; see Methods). The first four PCs define an orthonormal basis for the control vector **u** that is sufficient to capture 61% of the measured variation in the Fourier coefficients. We assumed that the four PCs could be controlled independently on each wing, and used them together with the wingbeat frequency *f* to define the 9 elements of the control vector **u**. It is an open question whether this approach is sufficient to describe all of the important kinematic variation in blowfly flight control, but our use of these four PCs is a pragmatic choice to ensure that the dimension of the resulting control input vector **u** is the same as that of the state vector **x**, resulting in a fully-actuated, rather than under- or over-actuated, system. Furthermore, in a qualitative sense, the first four PCs already capture the key kinematic control inputs that are known to be important in insect flight control (*43*), including coupled changes in stroke amplitude and stroke plane (Fig. 3C), and changes in the timing and extent of wing rotation at or around stroke reversal (Fig. 3B).

#### Physical modeling of the state equation

We used synchrotron-based X-ray microtomography (Fig. 3A) to estimate the inertia tensor of *Calliphora* for N=3 freshly-killed individuals, and used a Reynolds-averaged Navier-Stokes (RANS) solver (Fig. 3D) to model the aerodynamic forces and moments acting at the wing hinge under the reference kinematics. Because the flight dynamics model is linearized about equilibrium, we adjusted the assumed body mass, body drag, and wing hinge moment arm so that the aerodynamic forces and moments balanced the gravitational force when integrated over the reference wingbeat kinematics. We then ran a computational experiment (*44*) in which we simulated the aerodynamic effect of small perturbations to the insect’s motion state in **x**. These perturbations are aerodynamically equivalent to the gust disturbances in **d**, so having estimated the partial derivatives of the wingbeat-averaged aerodynamic forces and moments using a zero-intercept regression model (Fig. 3E), we were able to parameterize the system matrix **A** and disturbance matrix **G**. We parameterized the control matrix **B** in a similar manner for symmetric versus asymmetric control inputs, thereby completing our modeling of the state equation (Eq. 1).

### Co-tuning of Physics & Physiology

Having fully parameterized our state-space model of blowfly flight (Eqs. 1–2), it remains to identify the functional principle that underpins the physiological tuning of the LPTCs. We begin by examining the natural dynamics of the physical system, as a prerequisite for the more advanced control-theoretic analyses that follow.

#### Eigenstructure of the flight dynamics

The system matrix **A** has a similar eigenstructure (Fig. 3F) to most other models of insect flight dynamics (*18*), describing a characteristic set of symmetric versus asymmetric, stable versus unstable, and oscillatory versus monotonic motions. We summarise these by reporting the non-dimensional period 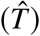 and/or time constant 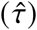 of each mode expressed relative to the insect’s wingbeat period. The symmetric modes are dominated by coupled pitch-heave motions, comprising a pair of fast 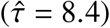 and slow 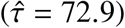 stable monotonic subsidence modes, and a slow unstable oscillatory mode 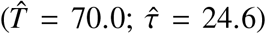. The asymmetric modes are dominated by coupled roll-yaw motions, comprising a slow but heavily-damped oscillatory mode 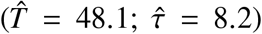, a fast stable monotonic subsidence mode 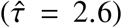, and a fast unstable monotonic divergence mode 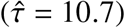. The time constants of the unstable modes are large enough that the instability they describe will develop on a timescale of tens of wingbeats in the absence of closed-loop control. As the period of each oscillatory mode is also an order of magnitude longer than the wingbeat period, these results validate our earlier assumption that the aerodynamic forces and moments may be replaced by their wingbeat averages when modeling the rigid-body flight dynamics (Eq. 1). Moreover, as blowfly flight is inherently unstable in respect of both symmetric and asymmetric motions, it follows that the insect must use the output of its sensors to command closed-loop flight stabilization. That being so, how has evolution tuned the visual physiology of the blowfly in relation to its flight dynamics?

#### Formalization of the mode-sensing hypothesis

The tuning of the LPTCs is characterised by the normalized row vectors of the bilateral output matrix **C**, each of which represents a specific direction of symmetric or asymmetric self-motion that the LPTCs are best-tuned to detect (*5*). The mode-sensing hypothesis predicts that these physiologically-preferred directions of self-motion should be matched to certain dynamically-significant directions of self-motion determined by the animal’s flight dynamics (*3*). For the unstable system described by Eqs. 1–2, those dynamically-significant directions are characterised by a set of real symmetric matrix structures called Gramians, which are defined in the frequency (*ω*) domain as:

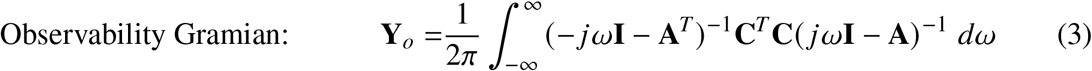

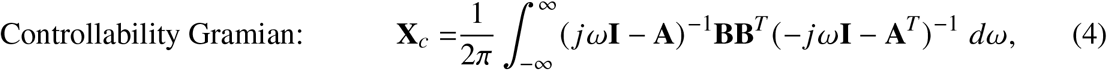

where **I** is the identity matrix (*45*). The disturbance-sensitivity Gramian **X**_*d*_ is composed similar to the controllability Gramian **X**_*c*_, replacing **B** with **G**. These Gramians are evaluated as solutions to a combined pair of Riccati and Lyapunov equations (see Supplementary Materials), but it is clear by inspection of Eqs. 3–4 that they relate to the interaction between the natural response of the system described by the system matrix **A**, and either the output matrix **C**, the control matrix **B**, or the disturbance matrix **G**. Each Gramian therefore relates to one of the distinct flows of signal energy summarised by the red arrows in Fig. 1.

To explain their dynamical significance more formally, we note that the orthonormal eigenvectors 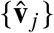 and ordered eigenvalues {*λ* _*j*_} of each Gramian define the principal axes of an *n*-dimensional ellipsoid with semi-axis lengths 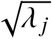, the longest axes of which represent the most dynamically-significant directions in the insect’s state space (*46*). For example, the eigenvectors and eigenvalues of the controllability Gramian **X**_*c*_ define the principal axes of the controllability ellipsoid 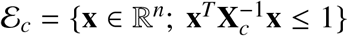. The dynamical significance of this structure can be seen by noting that the controllability Gramian **X**_*c*_ is defined (*45*) such that the quantity 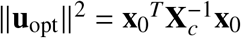 represents the minimum control input energy required to: (i) evolve the stable portion of the system to a given state **x** = **x**_0_; and (ii) regulate the unstable portion of the system to the origin **x** = **0**. It follows that the longest axes of the controllability ellipsoid ℰ_*c*_ encode the most-controllable directions in the insect’s state space, representing those specific self-motions that can be produced with the least input energy ∥**u**_opt_∥^2^ at the controls.

The observability and disturbance-sensitivity ellipsoids are constructed similarly, such that the observability ellipsoid ℰ_*o*_ encodes the most-observable directions (i.e., those self-motions that characteristically yield the most output energy at the sensors), and the disturbance-sensitivity ellipsoid ℰ_*d*_ encodes the most-sensitive directions (i.e. those self-motions that can be excited with the least input energy in a gust). These three ellipsoids thereby describe the specific self-motions that the insect is best-able to drive (ℰ_*c*_), best-equipped to estimate (ℰ_*o*_), and most-prone to experience (ℰ_*d*_). Aligning the sensors to any of these sets of dynamically-significant directions would therefore reflect a different optimization principle associated with signal energy flow through the system (Fig. 1). Comparing these dynamically-significant directions with the preferred directions of the LPTCs (Fig. 4B-D) allows a formal test of the mode-sensing hypothesis, and the identification of any underlying optimality principle within the control-theoretic framework of observability, controllability and disturbance-sensitivity (*47*).

#### Directional tuning of the LPTCs

We quantified the alignment of the preferred directions of the LPTCs to the dynamically-significant directions of self-motion by taking their absolute inner products 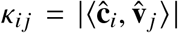. However, as the inner product of any symmetric-asymmetric pairing is identically zero, we use 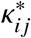 to distinguish symmetric-symmetric or asymmetric-asymmetric pairings for which 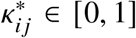. This analysis shows that the differenced responses of the three V-cells and VS2–10 are very strongly aligned 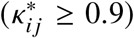 with the most-controllable direction of asymmetric motion, whilst the differenced response of VS1 is similarly strongly aligned with the second most-controllable direction of asymmetric motion (Fig. 4B). Likewise, the summed responses of the subset {V1, Vx, VS1-2, VS8–10} are strongly aligned 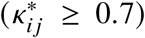 with the most-controllable direction of symmetric motion (Fig. 4B). The same holds true for the disturbance-sensitivity directions, but the summed and differenced responses of the vertical system cells are only weakly aligned with the most-observable directions of symmetric and asymmetric motion 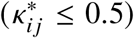. In contrast, the summed responses of the HS and H-cells are all strongly aligned 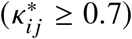 with the most-observable direction of symmetric motion (Fig. 4C), with those of the subset {HSE, H1, H2, Hx} being especially so 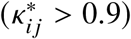.

#### Overall tuning of LPTC sub-populations

To assess the tuning of each sub-population of LPTCs to each set of dynamically-significant directions, we defined their weighted mean alignment (*η*) as:

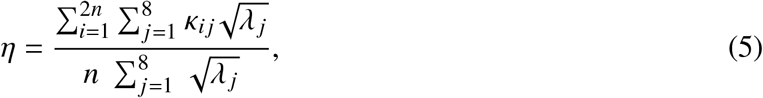

which measures the extent to which the preferred direction vectors of an entire sub-population of LPTCs encode the longest axes of a given ellipsoid, and generalizes to the case where the preferred directions of the LPTCs used to calculate the {*κ*_*ij*_} are replaced with a mirrored set of randomized direction vectors drawn from a uniform distribution on the unit sphere in ℝ^8^. A Monte Carlo analysis run over 100,000 such sets yields an expected weighted mean alignment of 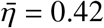 under the null hypothesis. Applying a Bonferroni correction to control the Type I error rate at *α* = 0.05 (Fig. 4D), we find that the VS-cells encode the most-controllable and most-sensitive directions much more strongly than expected by chance (*η* = 0.73 and *η* = 0.76, respectively; two-tailed *p* < .0025) and the most-observable directions more weakly (*η* = 0.17; two-tailed *p* < .0025). The V-cells display similar properties, also encoding the most-controllable and most-sensitive directions much more strongly than expected by chance (*η* = 0.75 and *η* = 0.80, respectively; two-tailed *p* < .0025). In contrast, the HS-cells do not encode any set of dynamically-significant directions any more strongly than expected by chance (*η* ≤ 0.42), and the H-cells encode the most-sensitive directions much more weakly (*η* = 0.07; two-tailed *p* < .0025). It follows that the VS- and V-cells are tuned to encode the effects of control inputs and gust disturbances, whereas the H-cells are tuned to observe characteristics of the optic flow field that are comparatively insensitive to disturbances.

#### Joint optimization of sensing & control

The properties of controllability, observability, and disturbance-sensitivity relate to signal energy flow to or from the system state, so depend upon our choice of coordinate system for the state vector **x**. That choice is meaningful for a technological system whose internal state is represented explicitly in its programming, but is ambiguous for a biological system whose internal state cannot be directly observed. This ambiguity is eliminated by the alternative hypothesis that instead of being tuned to optimize any one such property in a given state space, the preferred directions of the LPTCs are matched to the directions of self-motion that jointly optimize controllability and observability, or disturbance-sensitivity and observability. A system that implemented this principle would be globally optimal in the sense that it would maximize the transfer of signal energy from input to output, and hence unique in the sense that it would not depend on our choice of coordinate system for the state. For example, if evolution has tuned the vertical system cells to detect the effects of control inputs, then we should expect them to be strongly aligned to the joint most-controllable/observable directions. Conversely, if evolution has tuned the horizontal system cells to observe characteristics of the optic flow field that are robust to gust disturbances, then we should expect them to be weakly aligned to the joint most-sensitive/observable directions.

The jointly-optimized directions of self-motion are given by the normalized column vectors 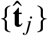 of the transformation matrix **T**^−1^, where 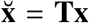 is a balancing transform that equalizes and simultaneously diagonalizes the transformed Gramians. Applying this balancing transform, which is unique up to multiplication by a sign matrix, we have either 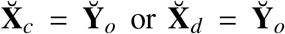, where 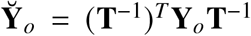 is a diagonal matrix. The diagonal entries of this matrix are the system’s Hankel singular values, which may be calculated directly as 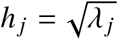 where {*λ*_*j*_} are the ordered eigenvalues of **Y**_0_**X**_*c*_ or **Y**_0_**X**_*d*_. The Hankel singular values are similarity invariants that do not depend on the choice of coordinate system for **x**, and they measure the degree of joint observability and controllability or disturbance-sensitivity in the directions 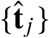 in the original coordinate system. Importantly, whilst the eigenvectors of the balanced Gramians in the new coordinates are orthogonal, the directions that they define in the original state space are not.

To assess the overall tuning of the LPTCs to these jointly-optimized directions, we calculated their absolute inner products as 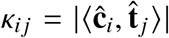, reporting their weighted mean *η* using the Hankel singular values 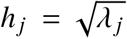 as the weights in Eq. 5. This analysis (Fig. 4D; see also Fig. 5A,B) confirms that the VS and V-cells encode the most-controllable/observable (*η* ≥ 0.75) and most-sensitive/observable directions (*η* ≥ 0.74) much more strongly than expected by chance (two-tailed *p* < .0025). The VS- and V-cells thereby embed the principle of encoding the directions in state space for which signal energy flow through the system is maximized. In other words, they are tuned to sense those modes of motion whose excitation yields the greatest sensor output for a given input of energy at the controls or in a gust. In contrast, the H-cells encode the most-sensitive/observable directions much more weakly than expected by chance (*η* = 0.13; two-tailed *p* < .0025). In other words, they are tuned to be insensitive to those modes of motion that are most readily excited by aerodynamic disturbances.

**Figure 5:**
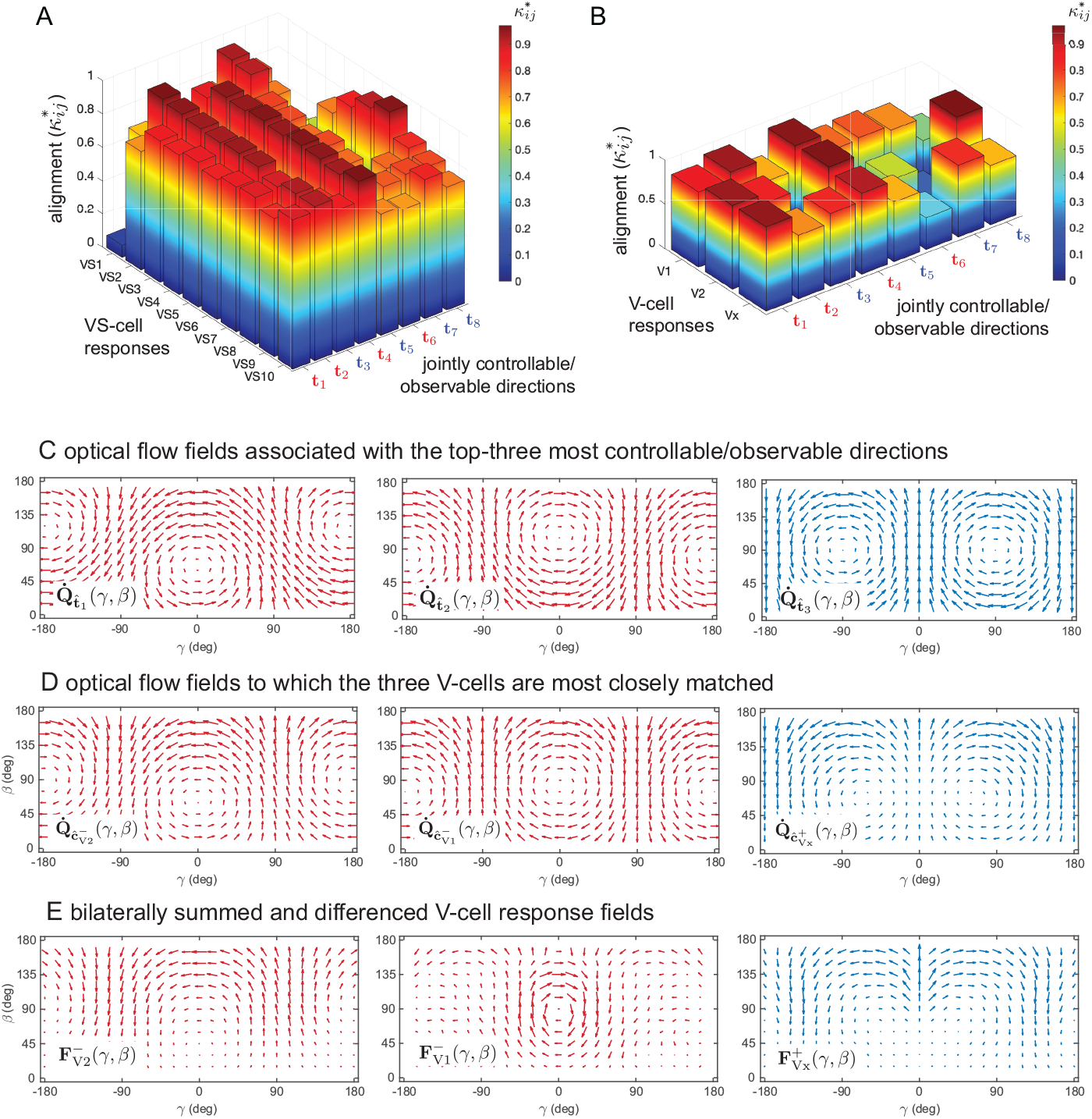
Tuning of the vertical system LPTCs to the fly’s jointly controllable/observable directions of self-motion. **(A,B)** Signal energy flow is maximized when a system’s sensors are matched to its most-controllable/observable directions of self-motion. To demonstrate the closeness of this tuning in the blowfly, we first quantified the alignment of the preferred directions of the LPTCs to the ordered symmetric (blue) or asymmetric (red) column vectors {**t**_1_, …, **t**_8_} of the inverse balancing transformation matrix. These vectors span the entire controllable/observable subspace of the insect, and the bilaterally summed and differenced response fields of the VS- and V-cell pairs are shown to be strongly aligned with them all. **(C-E)** To demonstrate this visually, we also computed the re-projected optic flow fields corresponding to: (C) the top three most controllable/observable directions of symmetric (blue) or asymmetric (red) self-motion; and (D) the preferred directions of symmetric (blue) or asymmetric (red) self-motion for the three V-cell pairs. Note the closeness of the match. **(E)** The summed and differenced response fields of the V-cell pairs also resemble the re-projected optic flow fields corresponding to the joint most-controllable/observable directions of motion, but do so less closely owing to spatial variation in the cells’ local motion sensitivity, which is expected to embed a nearness distribution corresponding to a natural environment (*16*).

#### A functional principle of visuomotor tuning

The preceding analyses compare the randomly-generated directions or preferred directions of the LPTCs against the biological ground truth of the parameterized state space model (Eqs. 1–2). This approach enables us to draw conclusions on the directional tuning of the individual LPTCs, but risks circular reasoning because their preferred directions also define the output matrix **C** that is used to generate the observability Gramian 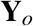 (Eq. 3). This circularity can be avoided altogether by composing a semi-random observability Gramian 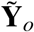 from the original system matrix **A** and a random output matrix 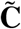 formed by generating 19 random direction vectors, mirroring these to yield 19 mirror-symmetric pairs, and taking their sums and differences to yield a bilateral output matrix with 38 rows. We then use 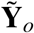 to compute the randomized Hankel singular values as 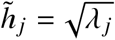, where the {*λ*_*j*_} are the ordered eigenvalues of 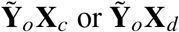.

Fig. 4E shows three measures of the Hankel singular values associated with the fly’s joint controllability/observability directions fall at the upper extreme of the null distribution of 100,000 randomly-generated sets. The first measure is the Hankel norm for the open loop system, or the largest Hankel singular value *h*_1_. The second is the Hankel-Frobenius norm,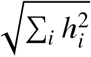, and the third is the Hankel Nuclear norm, Σ_*i*_ *h*_*i*_. This confirms our conclusion that the preferred directions of the blowfly motion vision system are specifically adapted to maximize the flow of signal energy from control inputs to sensor outputs, where signal energy is defined for an arbitrary vector signal **w**(*t*) as

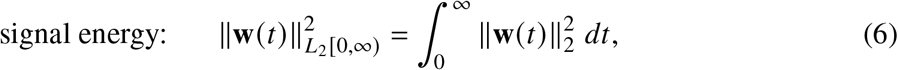

and ∥ · ∥_2_ is the Euclidean vector norm. Moreover, the first Hankel singular value provides an upper bound on signal energy flow from input to output, and is higher for the biological ground truth than for any of the randomly-generated systems in Fig. 4E. Similar conclusions hold for the fly’s joint disturbance-sensitivity/observability directions, and hence for the flow of signal energy from gust disturbances to sensor outputs (Fig. S4B,D). Finally, as the neck motor neurons that drive optokinetic head movements display similar response properties to the VS and HS cells (*33*), we infer that the same principle of maximizing signal energy transfer is also likely to apply at the level of the neck motor system driving any compensatory head movements.

#### Robustness of conclusions

To avoid making any assumptions on how the LPTC outputs are combined, we verified that the same conclusions hold when the Hankel singular values are calculated for the unilateral output matrix **C**′ as opposed to the bilateral output matrix **C** (Fig. S4). We also tested how the assumed nearness distribution of the visual environment influences the fly’s Hankel singular values, by synthesizing 100 perturbed output matrices and observability Gramians with respect to two different environmental configurations: a cuboid environment generated by perturbing the 2 m cube used in the analyses above, and a segmented ellipsoid environment generated by perturbing a sphere of equal nominal volume (Fig. S3A,C). The parameters defining each configuration were selected at random from a uniform distribution with 50% variation from their nominal values to generate variability and asymmetry in the assumed visual environment. The resulting distributions of perturbed Hankel singular values again remain at the extreme upper end of the null distribution (Fig. S3B,D). We conclude that the directional tuning of the LPTCs maximizes signal energy transfer between the inputs and outputs of the system, as opposed to maximizing conventional design criteria such as the accuracy of state estimation.

## Discussion

How does the LPTCs’ tuning embed the functional principle of maximizing signal energy transfer from input to output? As we have shown (Fig. 4), the vertical system LPTCs are strongly aligned to the insect’s most-controllable/observable directions of self-motion. But more than this, their alignment is high for all 8 of the joint controllability/observability directions — not just those which are the most controllable/observable (Fig. 5A,B). Specifically, every one of the joint controllability/observability directions is strongly aligned 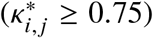 with at least one VS-cell and at least one V-cell (Fig. 5A,B), and the same result holds true for the joint disturbance-sensitivity/observability directions. In contrast, the horizontal system cells encode the joint controllability/observability directions much less strongly (Fig. 4D), effectively encoding a region of state space that is of lesser dynamical significance to the insect, and which is likely to be of greater significance in guidance and navigation tasks.

How is this combination of breadth and specificity possible? In principle, the 10 pairs of VS-cells have sufficient redundancy to encode any set of 8 directions strongly, but the same cannot be said of the 3 pairs of V-cells. The strength of the V-cells’ alignment to all of the joint controllability/observability directions instead reflects the fact that these 8 directions are highly non-orthogonal, describing a narrow region of the insect’s state space that is dominated by the same rotational motions as dominate the VS- and V-cell responses (Fig. 2B-E). Specifically, the natural modes of motion of a blowfly (Fig. 3F) are dominated by pitch-heave and roll-yaw dynamics (Table S6), which are the same self-motions that the vertical system LPTCs are tuned to sense. This is the region of state space that is of the greatest dynamical significance to the insect, and the hidden layer of V-cells embeds it in its entirety (Fig. 5B).

The strength of this embedding can be visualised by comparing the summed and differenced response fields of the V-cells (Fig. 5E) to the optic flow fields corresponding to their preferred directions of self-motion (Fig. 5D) and to the optic flow fields corresponding to the top three most-controllable/observable directions for the insect (Fig. 5C). The closeness of the match is striking, and this together with the broader correspondence between the vertical system LPTCs and the joint controllability/observability and disturbance-sensitivity/observability directions makes the VS- and V-cells well-suited to modulating flight stabilization/control. In contrast, the horizontal system LPTC responses are dominated by forward motion and yaw-sideslip. With the exception of yaw, these motions are of lesser dynamical significance, but they include the equilibrium forward-flight condition about which the dynamics are linearized. It follows that the horizontal system cells are better suited to encoding steady-state properties of the insect’s self-motion associated with its navigational state and guidance behaviors, which are also the directions of self-motion that are most robust to disturbance.

What are the functional benefits of structuring a system in this way? Intuitively, it makes sense to optimize a system’s sensors in relation to the endogenous inputs and exogenous disturbances that excite its motion, but how does this work in the context of closed-loop control? The transfer of signal energy from input to output is maximized by optimizing the storage and retrieval of signal energy to and from the system state. For an open-loop system, signal energy stored in the state **x** is released through the evolution of the natural modes of the system matrix **A**, which may be shaped arbitrarily through closed-loop control. Applying the principle of maximizing the Hankel singular values by tuning the sensing directions in **C** to the actuation directions of **B** through the natural modes of **A** balances the open-loop system so that it is optimized for maximum achievable closed-loop performance. This approach of tuning an open-loop system to maximize potential closed-loop performance, independent of the feedback architecture, is well documented and has significant precedent in the sensor placement and selection literature (*48*).

The evolutionary principle that we have identified of maximizing signal energy transfer from motor input to sensor output represents a radical departure from the design of current technological systems. In conventional engineering practice, sensor placement is usually optimized at a late stage of platform design, and typically aims to maximize the accuracy of state estimation. This is achieved by maximizing the signal-to-noise ratio at the sensors, which means placing them to optimize observability (*23, 49*). Tuning the LPTC response fields in this manner would yield sensor outputs with the best possible signal-to-noise ratio, but if the system was rarely excited in those directions by its own actions or by disturbances, then it would not be energetically efficient to encode them. In contrast, natural selection tends to produce neural architectures that prioritize energetic and hence computational efficiency (*50*), so it is reasonable to suppose that the principle of maximizing signal energy throughput might also extend to the sensorimotor systems of other organisms.

As the field of robotics transitions from platforms with sensorimotor architectures composed of small numbers of discrete sensors and actuators to architectures with continuum sensing and actuation, new design principles will be required. In order to achieve optimal performance, synthesis approaches that simultaneously consider the specification of sensors, actuators, and platform dynamics will be critical. Embodied design principles that have produced nature’s most effective and agile organisms (*51*), like the joint maximization of signal energy storage/retrieval uncovered here, have the potential to revolutionize the early stage design process and maximize the performance of future engineering systems. Such bio-informed design principles will prove especially relevant in applications that—like living organisms—are resource-constrained by computational capacity and power or energy density.

## Materials and Methods

### Electrophysiological characterisation of LPTC response fields

#### Animals and dissection method

Adult male and female blowflies (*Calliphora vicina*) were taken from a laboratory colony at Imperial College London where they were kept on a 12:12 hour light:dark cycle. Flies were dissected following a standardized procedure (*52*): after removing the legs, proboscis, and wings, the resulting wounds were sealed with beeswax before fixing the animal to a dedicated holder, with the thorax tilted 45° down relative to the head. The rear head capsule was opened using a micro-scalpel, and fat tissue, air sacs, and some tracheae were removed to enable placement of extracellular electrodes in the lobula plate. Saline solution (*53*) was added to keep the neural tissue moist. The centre of the head was positioned at the centre of a goniometric visual stimulation device, and aligned using the symmetrical deep pseudo-pupil method (*54*) at a precision of ±1° in head roll, pitch and yaw.

#### Animals and dissection method

Adult male and female blowflies (*Calliphora vicina*) were taken from a laboratory colony at Imperial College London where they were kept on a 12:12 hour light:dark cycle. Flies were dissected following a standardized procedure (*52*): after removing the legs, proboscis, and wings, the resulting wounds were sealed with beeswax before fixing the animal to a dedicated holder, with the thorax tilted 45° down relative to the head. The rear head capsule was opened using a micro-scalpel, and fat tissue, air sacs, and some tracheae were removed to enable placement of extracellular electrodes in the lobula plate. Saline solution (*53*) was added to keep the neural tissue moist. The centre of the head was positioned at the centre of a goniometric visual stimulation device, and aligned using the symmetrical deep pseudo-pupil method (*54*) at a precision of ±1° in head roll, pitch and yaw.

#### Extracellular recording and visual stimulation method

We used extracellular tungsten electrodes with 3 MΩ impedance (FHC Inc., Bowdoin, ME, USA; product code: UEWSHGSE3N1M) to record the neural activity of the V1, V2, and Vx heterolateral LPTCs. The electrodes were placed within different target areas depending on the recorded cell type using the tracheal branching patterns of the lobula plate as landmarks. Neuronal signals were amplified using a custom-built differential amplifier with a nominal gain of 10k, sampled and digitized at 20 kHz using a NI-DAQ board (USB-6211), and stored on the hard drive of a PC. Response fields were characterized only when the signal-to-noise ratio between recorded LPTC spikes and background noise was greater than 2:1, in which case a simple threshold-based method was sufficient to reliably detect time-stamped spikes of the recorded cell.

We used a custom-built automated goniometric recording platform to present a moving visual stimulus at any specified azimuth (*γ*) and elevation (*β*) in the fly’s retinal coordinates. An LCD monitor (AOC AGON AG251FZ) was placed 0.3 m in front of the animal, running at a refresh rate of 240 Hz. We presented square-wave visual gratings (minimum, maximum intensity: 0.28, 265.70 cd m^−2^; Michelson contrast: 0.9979) moving perpendicular to their orientation typically at 1 or 3 Hz temporal frequency behind a circular aperture subtending 24°. Experiments where different temporal frequencies between 0.3-3 Hz were applied did not affect the self-motion preferences of the cells. To assess a cell’s local preferred direction (LPD) and local motion sensitivity (LMS), the grating was moved in 8 different directions at a spacing of 45°. Each motion stimulus was presented for 1 s, followed by a brief period of 0.5 s during which a homogeneous screen was presented at the mean luminance level. In total, the motion stimulus was presented at 84 positions over both eyes, covering nearly the entire 4*π* visual field.

#### Local preferred directions and local motion sensitivities

At any given stimulus position we calculated the vector sum from the responses to the 8 different stimulus directions. Magnitude and direction of the resulting vector were taken to indicate the LMS and LPD, respectively. Those response parameters were plotted as a function of azimuth (*γ*) and elevation (*β*) in a cylindrical projection of the fly’s spherical visual field to reveal the recorded cell’s global response field properties (Fig. 2B-E). Within each response field, all vectors were normalized to the maximum response measured. To facilitate recognition of the global response field organization, the measured data in Fig. 2B-E (black vectors) are complemented by interpolated data (colored vectors). The data collected using this method are in line with those previously gathered using a local stimulus that changed its motion direction continuously (*33, 52, 53*). Although we presented the motion stimulus at 84 positions over both eyes, the response fields plotted for the heterolateral LPTCs in Fig. 2C-E show only the data obtained upon visual stimulation within the visual hemisphere that results in the highest motion sensitivity (i.e., strongest directional-selective response), which is typically the visual hemisphere ipsilateral to the dendritic input region of the recorded cell.

### modeling of LPTC response properties

#### Preferred self-motion parameters

We estimated each cell’s preferred self-motion parameters using an iterative least squares algorithm (KvD) proposed by Koenderink and van Doorn (*55*). The KvD algorithm is applied to retrieve the angular velocity ***ω***, translational velocity **v**, and the local distance distribution *d* of self-motion that induces an optic flow field that best fits the global response field organization of the studied cell. Hence, if we consider a given LPTC to act as a matched-filter for optic flow (*16*), the KvD algorithm enables us to estimate which self-motion components of a moving fly would most strongly stimulate the cell. We applied a slightly modified version of the KvD algorithm in which we assumed a homogeneous distance distribution to obtain the preferred rotation axes of the V1, V2, and Vx-cells (Fig. 2G); the preferred rotation axes of the VS-cells (Fig. 2F) were computed using a similar method by (*26*). There were no significant sex differences in the preferred rotation axes of the three heterolateral LPTCs, and we therefore pooled the response field data across sexes.

#### Encoding of motion state by the LPTCs

The matched filter hypothesis proposed in (*16*) suggests that each LPTC’s output can be considered as a comparison between the cell’s response field and the optic flow fields generated during the animal’s self-motion, where each cell is tuned to sense a specific flow field and hence some specific combination of rotational and translational self-motion. Mathematically, this comparison can be modeled as an inner product on a discrete (*16, 56, 57*) or continuous (*58–60*) spatial domain. Here we define a spatial inner product between the instantaneous pattern of optic flow 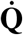 and a given tangential cell’s response field **F**_*i*_ as:

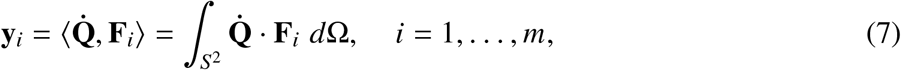

where *m* is the number of LPTC response fields under consideration.

The optic flow pattern 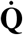 is the vector field of the relative velocity of visual contrast from objects in the environment projected into the tangent space *T*_**r**_*S*^2^ of the imaging surface (Fig. S1A). Its global structure depends primarily on the observer’s rotational velocity ***ω*** and translational velocity **v**. The translational contribution at each viewing direction **r** = (*γ, β*) is inversely scaled by the distance *d* (*γ, β*) from the imaging surface to the nearest object in the environment (Fig. S1B). Since *d* can be unbounded, the nearness function *μ*(*γ, β*) = 1/*d*(*γ, β*) is commonly used in the formulation. The instantaneous optic flow pattern 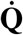 on a spherical imaging surface *S*^2^ for an arbitrary distribution of objects can be expressed as (*55*):

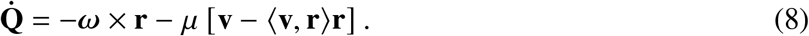

In order to formulate the optic flow pattern in closed form for the calculation of the spatial inner product 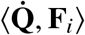, the shape of the nearness function and its dependence on the relative pose variables **q** = {*x, y, z, ϕ*_*s*_, *θ*_*s*_, *ψ*_*s*_} of the animal needs to be specified. Here {*x, y, z*} are the coordinates of the vantage point with respect to the inertial frame 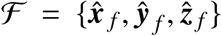 and {*ψ*_*s*_, *θ*_*s*_, *ϕ*_*s*_} are the 3-2-1 Euler angles of the stability frame 𝒮 relative to the inertial frame ℱ. For the analysis here, two classes of environment structure were considered, which included an enclosed rectangular prism and a segmented ellipsoid (Fig. S4). For each of these cases, the nearness *μ*(*γ, β*, **q**) is a piecewise-continuous function and the methodology for their derivation can be found in (*60*).

#### Formulation of the output matrix

For a given set of unilateral response fields {**F**_*i*_}, the collection of LPTC outputs form a nonlinear output equation **y** = **h**(**x**). To characterize the rigid body state information encoded by the selected set of measured response fields, each output is linearized about the reference flight equilibrium 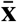. The resulting matrix entries in the unilateral output equation **y** = **C**′**x** (Eq. 2) are given by

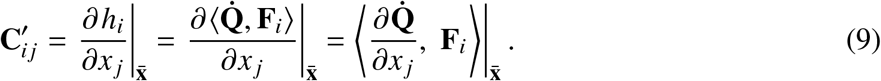

In this form, each row of the associated **C**′ matrix represents the state information present in the signal from a specific LPTC response field: that is, the direction it encodes in state space.

To develop the output matrix for the measured set of 19 left and right LPTC cell response fields, the raw data was first converted into stability frame coordinates according to the axis definitions of Fig. S2A. Note that (*γ, β*) = (0°, 90°) in the plotted response fields (Fig. 2B-E, 4A) corresponds to the ray line along the 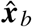 axis in Fig. S1B. The recorded LPD and LMS distributions within the response fields were smoothed with a 2D Gaussian filter and then approximated with up to 8th degree spherical harmonics in the azimuth and elevation directions to facilitate an accurate numerical spatial integration. For our baseline calculation, an enclosed rectangular prism environment (Fig. S4A) was assumed with scaling parameters (*g*_*N*_, *g*_*S*_, *a*_*E*_, *a*_*W*_, *h*_*D*_, *h*_*U*_) all set to a distance of 1 m, and as before the reference flight condition was set to the mean forward flight speed of *u*_*s*_ = 0.8509 m s^-1^ that we had measured (see below), such that 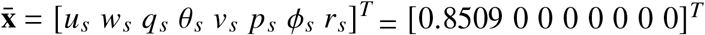 in the stability axes. The corresponding patterns of partial derivatives of the optic flow 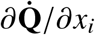 were computed by systematically perturbing each of the rigid body states and taking finite differences.

The results were used to compute the inner products numerically, resulting in the estimated response field state encoding shown in Table S7. To perform the subsequent analysis, we extract the entries according to the rigid body state vector from the flight dynamics model formulation, **x** = [*δu*_*s*_ *δw*_*s*_ *δq*_*s*_ *δθ*_*s*_ *δv*_*s*_ *δ p*_*s*_ *δϕ*_*s*_ *δr*_*s*_]^*T*^. The states {*δx*_*s*_, *δy*_*s*_, *δz*_*s*_, *δψ*_*s*_} typically are not included in the linearized dynamics since a homogeneous atmosphere assumption is employed. In the resulting matrix 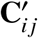, consecutive odd and even indices {*i* = 1, …, 38} correspond to the left and right cells of an LPTC pair. Finally, because the flight dynamics model (Eq. 1) splits into symmetric and asymmetric parts, we form the 38 normalized rows 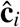 of the bilateral output matrix **C** (Eq. 2) by summing or differencing the responses of the *k* = 1, …, 19 mirror-symmetric pairs of LPTCs such that 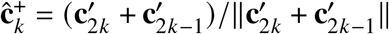 and 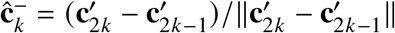 as shown in Fig. 4A.

#### Re-projected optic flow fields

Any given direction of self-motion will produce a specific optic flow field in a given visual environment. We re-projected the optic flow fields 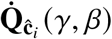 and 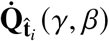 corresponding the specific self-motion directions 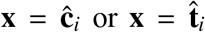 in state space (Fig. 5C-E) by substituting the expressions of the vectors ***ω*** = [*p q r*]^*T*^, **v** = [*u v w*]^*T*^, and **r** = [cos *γ* sin *β* sin *γ* sin *β* cos *β*]^*T*^ in the stability frame 𝒮, along with an analytical expression for the nearness of the environment *μ*(*γ, β*, **q**) into the representation for 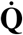 above. The resulting azimuth *γ* and elevation *β* components (Fig. S1) of 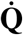 are given by

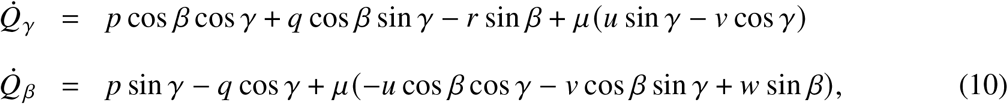

as plotted in Fig. 5C-E.

### Measurement and modeling of free-flight kinematics

#### Animals and experimental protocol

Larval *C. vicina* were reared on red meat at 20°C until pupation; the adult flies were fed on a combination of milk powder formula and mashed banana, and were flown from 2–3 days post-eclosion. Individuals were allowed to fly freely within a 1 m diameter opaque acrylic sphere with diffuse overhead lighting. The interior of the sphere was decorated with pieces of card to provide visual contrast, and an ultraviolet light was placed at the top to stimulate loitering flight maneuvers. High-speed video sequences were captured using four synchronized Photron SA3 cameras (Photron Ltd, West Wycombe, UK) with 180 mm macro lenses (Sigma Imaging Ltd, Welwyn Garden City, UK) viewing the insect through clear portholes in the upper hemisphere, recording at 3,800 fps and 768×640 pixels. Backlighting was provided by four infrared LED lights (Dragon1IR PowerStars LED, Intelligent LED Solutions, Thatcham, UK) operating at wavelengths well beyond the visible spectrum of the insect (*61*) (centroid wavelength: 850 nm; spectral bandwidth at 50% intensity: 30 nm full width at half maximum). Recordings were triggered as the insect passed through the centre of the sphere, capturing forward flight punctuated by fast saccadic maneuvers. In total, we recorded *N* = 2, 708 wingbeats from 205 maneuvering flights made by 28 individual blowflies, involving a broad range of wing kinematics including symmetric and asymmetric variation in stroke amplitude, stroke plane angle, and wing twist.

#### Kinematic reconstruction

The cameras were calibrated using a nonlinear least squares bundle adjustment routine (*62*) in Matlab (The Mathworks, Inc., Natick, MA), using images of a calibration grid presented in a wide range of positions and orientations. For the main analysis, we used background subtraction and automatic thresholding to segment the pixels, and used a shape-carving method to identify the set of voxels containing the wings and body (*41*). We reprojected the wing voxels as a mask for tracing the outline of the wing in each frame, and used the shape-carving algorithm on this linear feature to reconstruct the leading and trailing edges in three dimensions. We used the major axis of the body voxels to define the insect’s *x*-axis, and the line connecting the wing roots to define the insect’s transverse *y*-axis. We constructed a right-handed body axis system in which to measure the kinematics of the right wing, and a left-handed body axis system in which to measure the kinematics of the left wing. These were measured by defining an extrinsic *y*-*x*-*z* rotation sequence bringing the *x*-axis of a set of rotating axes initially aligned with the body axes into alignment with the wing chord connecting the trailing edge to the leading edge at some given spanwise position (*r*). The local pitch angle of the wing (*ω*), is defined as the first angle in this rotation sequence, and was measured at 6 evenly-spaced spanwise stations on the interval *r* ∈ [0.3, 0.8], where *r* is expressed as a proportion of wing length. We summarised the instantaneous spanwise variation in *ω* by fitting the regression model *ω*[*r*] = *ω*_0_ + *ω*_*r*_*r* + *ϵ* [*r*], where *ϵ* [*r*] is a Gaussian error term. We call *ω*_0_ the twist offset, and *ω*_*r*_ the twist gradient. The deviation angle *θ*_*w*_ and stroke angle *ϕ*_*w*_ represent the second and third angles in the extrinsic *y*-*x*-*z* rotation sequence, and describe the elevation and azimuth of the wingtip in a set of body axes originating at the wing root. It follows that the insect’s wing kinematics are measured by estimating *ϕ*_*w*_ [*t*], *θ*_*w*_ [*t*], *ω*_0_[*t*], *ω*_*r*_ [*t*] for the right and left wings separately at every sample time *t*.

#### Fourier series representations of wing kinematics

For each flight sequence, we fitted quintic smoothing splines modeling *ϕ*_*w*_ [*t*], *θ*_*w*_ [*t*], *ω*_0_[*t*], and *ω*_*r*_ [*t*] for each wing as analytical functions of continuous time *t*. The spline tolerance that we used for each kinematic variable was chosen to preserve information up to the 3^rd^ harmonic of wingbeat frequency for *ϕ*_*w*_ and *θ*_*w*_, and up to the 5^th^ harmonic for *ω*_0_ and *ω*_*r*_. We then used a piecewise linear transform to map continuous time *t* onto wingbeat phase *φ*(*t*), by taking the turning point of the summed angular velocity of both wingtips in the stroke plane to define *φ* = 0 as the start of the downstroke. Finally, we evaluated the splines at 101 evenly-spaced phases of each wingbeat on the interval *φ* ∈ [0, 2*π*], so that all wingbeats were directly comparable despite variability in the wingbeat period. Fitting each wingbeat separately, we used multiple regression with time-linear and time-periodic predictor variables to model the four primary kinematic variables *ϕ*_*w*_ [*φ*], *θ*_*w*_ [*φ*], *ω*_0_[*φ*], and *ω*_*r*_ [*φ*] as de-trended Fourier series of the form:

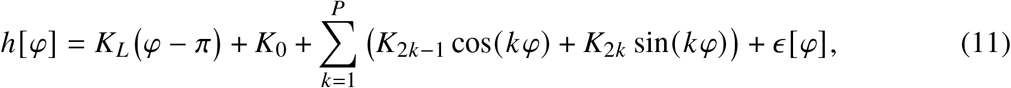

where *ϵ* [*φ*] is a Gaussian error term, and where *K*_*L*_ and *K*_0_ … *K*_2*P*_ are fitted coefficients. The time-linear coefficient *K*_*L*_ accounts for the fact that any actual wingbeat cycle is unlikely to begin and end in exactly the same kinematic state, and multiplies *φ* − *π* rather than *φ* so that this term has zero mean over the wingbeat cycle. The Fourier coefficients *K*_0_ … *K*_2*P*_ are fitted with *P* = 3 for *ϕ*_*w*_ and *θ*_*w*_, and with *P* = 5 for *ω*_0_ and *ω*_*r*_, to capture all of the harmonic content preserved by the quintic smoothing splines.

#### Functional principal components analysis

Collecting the Fourier coefficients for a single wingbeat together as 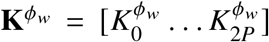 for the stroke angle *ϕ*_*w*_, and similarly for the other primary kinematic variables, we may summarise the time-periodic variation for all *N* wingbeat pairs in the matrix:

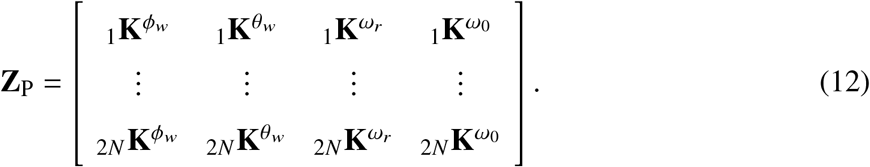

We used functional principal components analysis to decompose this matrix into a new set of time-periodic basis functions characterising the key kinematic couplings available for flight control. This was implemented by subtracting the column means from the matrix of Fourier coefficients **Z**_P_ to yield the centered matrix 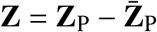, and computing its singular value decomposition:

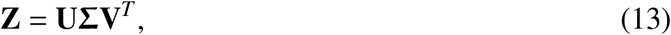

where **Σ** is a diagonal matrix containing the singular values of **Z**, which are the positive square roots of the eigenvalues of **Z**^*T*^ **Z** arranged in descending order. The columns of **V** contain the right-singular vectors of **Z**, which are the eigenvectors of **Z**^*T*^ **Z**, normalized such that **V**^*T*^ **V** = **I**. Because **Z**^*T*^ **Z** is a scalar multiple of the covariance matrix of **Z**, the orthonormal basis **V** that its eigenvectors define is aligned with the principal directions of the kinematic variation between wingbeats. Because each row of the principal component (PC) matrix **V** corresponds to one of the Fourier coefficients, each of its columns defines a distinct time-periodic kinematic coupling, which we refer to as PC1, PC2, etc.

### Aerodynamic modeling of stability and control derivatives

#### Computational fluid dynamics modeling

We performed three-dimensional Reynolds-averaged Navier-Stokes (RANS) simulations of the aerodynamics of the right wing of *Calliphora* using the OVERTURNS solver (*63, 64*). We simulated a reference condition of level symmetric forward flight with a freestream velocity of *U*_∞_ = 0.8509 m s^-1^ and a wingbeat frequency of *f* = 166.188 Hz, corresponding to the mean values measured for these variables over the 2,708 wingbeats whose kinematics we had recorded (see above). We used a mean body pitch angle *θ*_*e*_ = 30.365°, and defined a set of reference wing kinematics by taking the mean of **Z**_P_ over three consecutive wingbeats centered on the wingbeat most nearly achieving level flight among the subset of wingbeats for which the body acceleration was near-zero (< 0.5 m s^-2^). We assumed a pure laminar boundary layer based on the measured mean chord length of 2.7 mm and chord Reynolds number of 1,746. We used 720 time steps per wingbeat cycle with 12 sub-iterations, which allowed us to capture the unsteady flow characteristics with reasonable computational time. Tables S2 and S3 summarize the simulation parameters and input setup. The dimensions of the structured wing mesh were 195 (chordwise), 201 (spanwise), and 76 (wall-normal), and the initial wall normal spacing was 1 ×10^−5^ mean chord. Combined with the background mesh, this gave a total of 1.12 × 10^7^ node points.

#### Computational experiments

We perturbed the reference kinematics to simulate the aerodynamic effect of control inputs and small perturbations to the motion state of the body. We used the three components of translational velocity {*u, v, w*} and angular velocity {*p, q, r*} referred to the body axes B as the perturbed states, and used the wingbeat frequency *f* together with principal components PC1–4 of the time-periodic wingbeat kinematics as control perturbations. Among the six perturbed states variables {*u, v, w, p, q, r*}, the perturbation level was set at ±10% and ±20% of freestream velocity *U*_∞_ for the three flight velocities {*u, v, w*}, and at ±2.5% and ±5% of wingbeat frequency *f* for the three angular rates {*p, q, r*}. Among the five control variables, we perturbed PC1–4 by ±0.5 and ±1.0 standard deviations, and perturbed the wingbeat frequency *f* by ±1.5% and ±3%, corresponding to ±2.5 and ±5.0 Hz respectively. We computed the three components of aerodynamic force {*X, Y, Z*} and moment {*L, M, N*} referred to the body axes B (Fig. S2A). The time-averaged forces and moments had already converged reasonably by the start of the 3rd wingbeat cycle, so we obtained their wingbeat-averaged values by taking the mean of the forces and moments over the 3rd and 4th wingbeat cycles of each simulation.

#### Aerodynamic derivatives

In order to estimate the stability and control derivatives for our linearized flight dynamics model, we first subtracted the total aerodynamic forces and moments in each perturbed condition from those obtained in the reference flight condition, then fitted a zero-intercept linear regression through the origin to estimate the corresponding partial derivative of the aerodynamic forces and moments (Fig. 3B; Fig. S2D-E). The resulting aerodynamic derivatives are defined for the right wing using forces and moments resolved at the right wing hinge, and were mirrored to model the aerodynamic forces and moments on the left wing under the equivalent kinematics. The results for the left and right wings were then combined to estimate the stability derivatives (Table S4) and control derivatives (Table S5) resolved at the centre of mass, in a model enforcing equilibrium under the reference wing kinematics (see Supplementary Information for details). We used the unilateral control inputs to form sets of symmetric control inputs {*u*_0_, *u*_1_, *u*_2_, *u*_3_, *u*_4_} and asymmetric control inputs 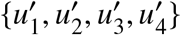 through symmetric changes in wingbeat frequency *f* and symmetric or asymmetric application of perturbations in PC1–4.

### Microtomographic estimation of inertia tensor

#### Animals and tomographic method

Two male and one female *C. vicina* from the Imperial College London colony were sealed in 0.5 mL Eppendorf tubes, having been fed and watered to satiation and then weighed. The tubes were placed individually in the TOMCAT beamline of the Swiss Light Source synchrotron facility and were irradiated at 12 keV beam energy until the flies were dead and no further motion artefacts occurred. A 100 *μ*m thick, Ce-doped LuAG scintillator was placed at a distance of 272 mm behind the sample to convert the transmitted X-rays into visible light. The resulting image was magnified 2-fold using an Edge 5.5 Microscope, and 1501 projection images were collected at 60 ms exposure time whilst the sample was rotated through 180°. To image the entirety of each fly, it was necessary to image three overlapping volumes in this way. Phase retrieval was performed using the Paganin algorithm (*65*), setting the real and imaginary parts of the deviation from one of the complex refractive index of the material to 1 × 10^−7^ and 1 × 10^−9^ respectively. Tomographic reconstruction was performed using a Fourier transform-based algorithm, resulting in voxels with an isotropic spacing of 3.25 *μ*m.

#### Estimation of inertia tensor

We segmented the tomograms automatically, using thresholding and morphological operations to mask voxels corresponding to the cross-section of the Eppendorf tube and any exterior voxels. The tomograms were then combined into one continuous stack across the three samples for each specimen, using unique cross-sectional features such as hairs to align the image stacks manually along their common longitudinal axis. The complete image stack was loaded into Fiji (*66*), and the BoneJ plugin (*67*) was used to calculate the mass moments of inertia about the principal axes of the specimen, assuming a uniform density of 1.1 g cm^−3^ appropriate to insect cuticle and muscle. The mass (*m*) estimates arrived at using this method (Table S1) were identical to the weights taken at the start of the experiment to within the 0.01 g readability of the balance. Because the moments of inertia about the first and second principal axes were identical to within ±1% (Table S1), we set *I*_*y*_ = *I*_*z*_ for the purposes of the flight dynamics modeling, and set *I*_*x*_ equal to the moment of inertia about the third principal axis. Assuming isometry, we non-dimensionalized the moments of inertia by dividing through by *m*^5/3^, then rescaled these using the value of *m* assumed in the flight dynamics model.

## Acknowledgments

The authors thank Zoe Turin, Gregory Gremillion, and Jishnu Keshavan for discussions related to the dynamical systems formulation. We acknowledge the Paul Scherrer Institut, Villigen, Switzerland for provision of synchrotron radiation beamtime at the TOMCAT beamline X02DA of the Swiss Light Source (SLS), and thank Christian Schlepütz for assistance with synchrotron data collection. We thank Ric Wehling, Johnny Evers, Pat Bradshaw, and Nick Rummelt for their longstanding support of this research and for many insightful conversations. We thank Henry Cerbone, Alex Yarger, Ben Campbell, and Jack Supple for helpful comments.

## Author contributions

J.S.H., H.G.K, and G.K.T. conceived the presented idea with R.W.Z. H.G.K conceived and supervised the electrophysiological experiments: J.V.H. developed the electrophysiology setup, C.R. and Y.Y, carried out the recordings, and analysed the electrophysiological data. G.K.T. and S.M.W. conceived and supervised the wing kinematic measurements: I.L.D. and S.M.W. carried out the videogrammetric experiments and analysis, and I.N. and G.K.T. carried out the functional principal components analysis. G.K.T. conducted the synchrotron experiments with

B.M. and analysed the tomographic data. J.B. supervised the CFD simulations: C.B., Y.S.J, and C.S. created and performed the simulations, and J.S.H., Y.S.J, and G.K.T carried out the data analysis.

J.S.H. and G.K.T. developed the theoretical framework: J.S.H., A.H., and G.K.T. conceived and performed the dynamical systems analysis. J.S.H. and G.K.T. wrote the manuscript with H.G.K. All authors participated in reviewing the manuscript.

## Data and materials availability

The data and code that support the findings of this study will be made available in figshare with the identifier doi:10.6084/m9.figshare.25504879. Correspondence and requests for materials should be addressed to G.K.T., J.S.H., and H.G.K.

## Supplementary materials

### Flight dynamics modeling

As outlined in the main text, we have parameterized a linear time-invariant (LTI) flight dynamics model:

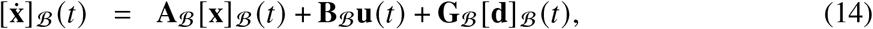

where the state vector [**x**]_B_ = [*δu δw δq δθ δv δ p δϕ δr*]^*T*^ is sufficient to describe the 6-DoF rigid body motions of the insect insofar as these motions influence the wingbeat-averaged aerodynamic forces {*X, Y, Z* } and moments {*L, M, N*} resolved in the principal axes of the body 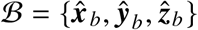 as described in Fig. S2A. Here, *δ* denotes a small perturbation from equilibrium, where {*u, v, w*} are the components of translational velocity along the principal axes of the body, where { *p, q, r* } are the components of angular velocity about these body axes, and where {*θ, ϕ*} are the pitch and bank angles of the insect. These Euler angles are defined as the second and third rotations in an intrinsic 3-2-1 rotation sequence bringing the body axes to their final orientation, starting from an initial configuration in which the *z*-axis is vertical. Note that the first Euler angle describing the azimuth *ψ* about the body *z*-axis is not included in the system state, because it has no influence on the flight physics.

The linearization of the equations of motion for a rigid flying body using small perturbation theory is dealt with in most flight dynamics texts, so is not repeated here. In brief, we modeled the insect as a symmetric rigid body subject to wingbeat-averaged aerodynamic forces and moments. These are described as linear functions of the motion state variables {*u, v, w, p, q, r*} and longitudi-nal and lateral control input variables **u**_long_ = [*u*_0_ *u*_1_ *u*_2_ *u*_3_ *u*_4_]^*T*^ and 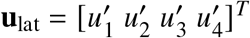 for small perturbations from a symmetric rectilinear equilibrium flight condition. See (*17, 18, 68*) for discussion of this approach in the context of insect flight dynamics, including the wingbeat-averaging of the aerodynamic forces and moments, which is an approach borrowed from simplified helicopter flight dynamics modeling. In the following, we assume that the relevant quantities are all expressed in the body frame ℬ, and therefore drop the coordinate frame notation ℬ hereafter.

With these definitions and assumptions, we model the longitudinal flight dynamics 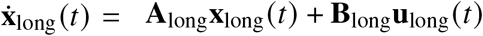 as

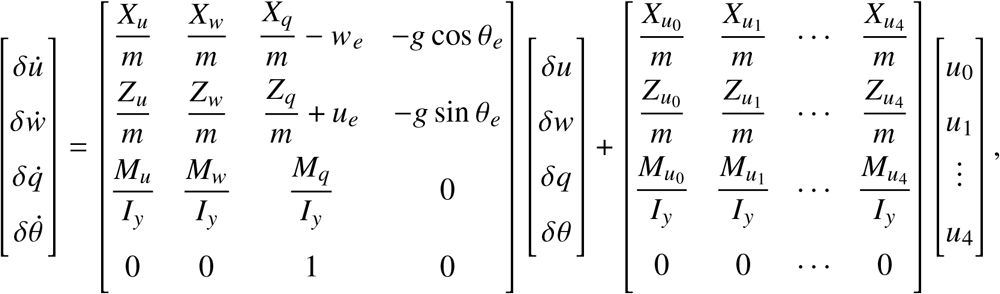

where {*u*_*e*_, *w*_*e*_, *θ*_*e*_ } are the values of {*u, w, θ*} at equilibrium, where *g* is gravitational acceleration, *m* is body mass, and *I*_*y*_ is the moment of inertia about the body *y*-axis. Here the control input *u*_0_ denotes a change of wingbeat frequency with respect to equilibrium, whilst the remaining control inputs {*u*_1_, · · ·, *u*_4_} denote mirror-symmetric application of the time-periodic kinematic couplings in the directions defined by PC1 through to PC4 (see Methods). Quantities of the form *X*_*u*_, *Z*_*w*_, *M*_*q*_, etc. are stability derivatives, denoting the partial derivatives ∂*X*/∂*u*, ∂*Z*/∂*w*, ∂*M*/∂*q*, etc. Likewise, quantities of the form 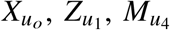,etc., are control derivatives, denoting the partial derivatives ∂*X*/∂*u*_0_, ∂ *Z*/∂*u*_1_, ∂*M*/∂*u*_4_, etc. Derivatives were estimated by regressing the wingbeat-averaged forces and moments predicted using CFD on the relevant state variables and control input variables (see Methods). Note that for the purposes of the flight dynamics modeling described here, we regress the perturbed forces and moments on the perturbed state variables and control input variables, and therefore force the regressions through the origin.

If we assume that external gusts **d**(*t*) = {*u*_*d*_, *v*_*d*_, *w*_*d*_, *p*_*d*_, *q*_*d*_, *r*_*d*_ } are acting on the animal, then we may replace the motion state variables {*u, v, w, p, q, r* } used to calculate the aerodynamic forces and moments (Δ*X*, Δ*M*, etc.) with an inertial perturbation and a disturbance term, such that *δu*_*a*_ = *δu* − *u*_*d*_, *δr*_*a*_ = *δr* − *r*_*d*_, etc. Under this assumption, the forces and moments on the insect are now a function of both the inertial and the gust components. For instance,

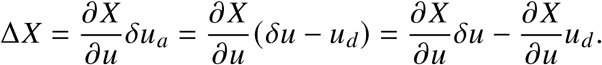

Therefore the way the gust *u*_*d*_ influences the dynamics 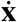 is through the term −∂*X*/∂*u*, which is the negative of the aerodynamic portions of the **A** matrix. The longitudinal flight dynamics model with gusts are then modeled as 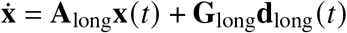 with

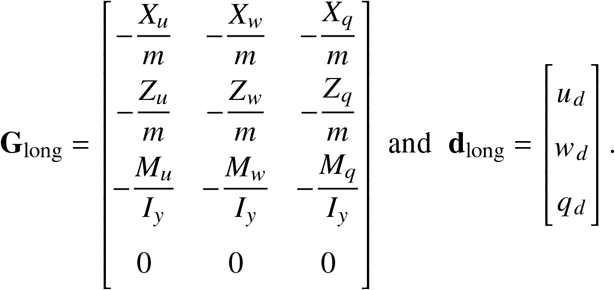

The lateral flight dynamics 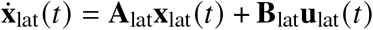 are modeled as

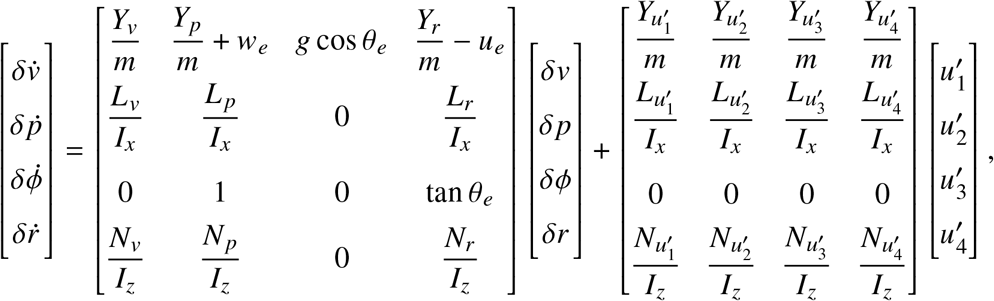

where the notation is similar, save that the control inputs 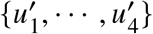 denote anti-symmetric application of the time-periodic kinematic couplings in the directions defined by PC1 through to PC4 (see Methods). The lateral flight dynamics model with gusts 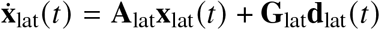 is derived in the same way as its longitudinal counterpart, with

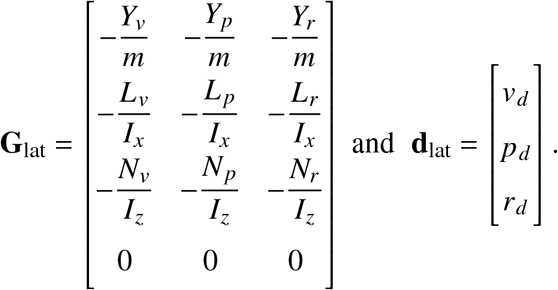

Note that these equations of motion are linearized about a symmetric rectilinear equilibrium flight condition with *v*_*e*_ = *p*_*e*_ = *q*_*e*_ = *r*_*e*_ = *ϕ*_*e*_ = 0, and neglect any gyroscopic forces on the beating wings. They further assume that the fastest natural modes of the system have a characteristic timescale much longer than the period of the wingbeat, such that there is no significant coupling between the time-periodic oscillations of the aerodynamic forces and the body’s motion. It is important to note that their simplified form reflects the fact that all of the forces and moments are resolved with respect to the principal axes of the insect. Finally, although we do not derive the LTI model of the flight dynamics explicitly above, another key feature of this model is that it is linearized about a state of equilibrium, which is a necessary condition for time-invariant equations of motion to result from the linearization.

Whilst it is reasonable to assume that the reference flight condition associated with the mean wingbeat in the functional principal components analysis will be close to equilibrium, there is no reason to expect that it will be exactly so. To guarantee the internal consistency of the model, we therefore enforce equilibrium, subject to the assumption: (i) that *v*_*e*_ = *p*_*e*_ = *q*_*e*_ = *r*_*e*_ = *ϕ*_*e*_ = 0 on grounds of symmetry; and (ii) that {*u*_*e*_, *w*_*e*_, *θ*_*e*_ } are each equal to the reference conditions assumed in the CFD modeling. We achieve this by making three assumptions. First, we assume that the magnitude of the vertical aerodynamic force predicted by the CFD under the reference flight conditions is exactly balanced by the insect’s body weight. We therefore solve for the value of body mass at which this assumption holds and set *m* equal to this value (5.7686 × 10^−5^ kg) in the flight dynamics model. Second, we assume that the net thrust predicted by the CFD under the reference flight conditions is exactly balanced by body drag in forward flight. This body drag does not feature explicitly in the flight dynamics model, because it is implicit in the treatment of the reference flight conditions from the CFD as an equilibrium condition. Third, noting that the aerodynamic moments predicted by the CFD are resolved at the root of the wing, we assume that any non-zero pitching moment about the wing root is balanced at the centre of mass by the vertical aerodynamic force. We therefore solve for the horizontal lever arm in the *xz*-plane for which this assumption holds and assume that the wing root is located on a known transverse lever arm along the *y*-axis that we measure directly from the tomograms (see Methods). Finally, we resolve all of the aerodynamic moments at the centre of mass prior to taking the stability and control derivatives above.

With these assumptions and parameterizations, the longitudinal and lateral **A, B**, and **G** matrices for a freestream velocity of *U*_∞_ = 0.8509 m/s and a body pitch angle *θ*_*e*_ = 30.365° expressed in the body frame ℬ for *Calliphora vicina* are as follows, in S.I. units:

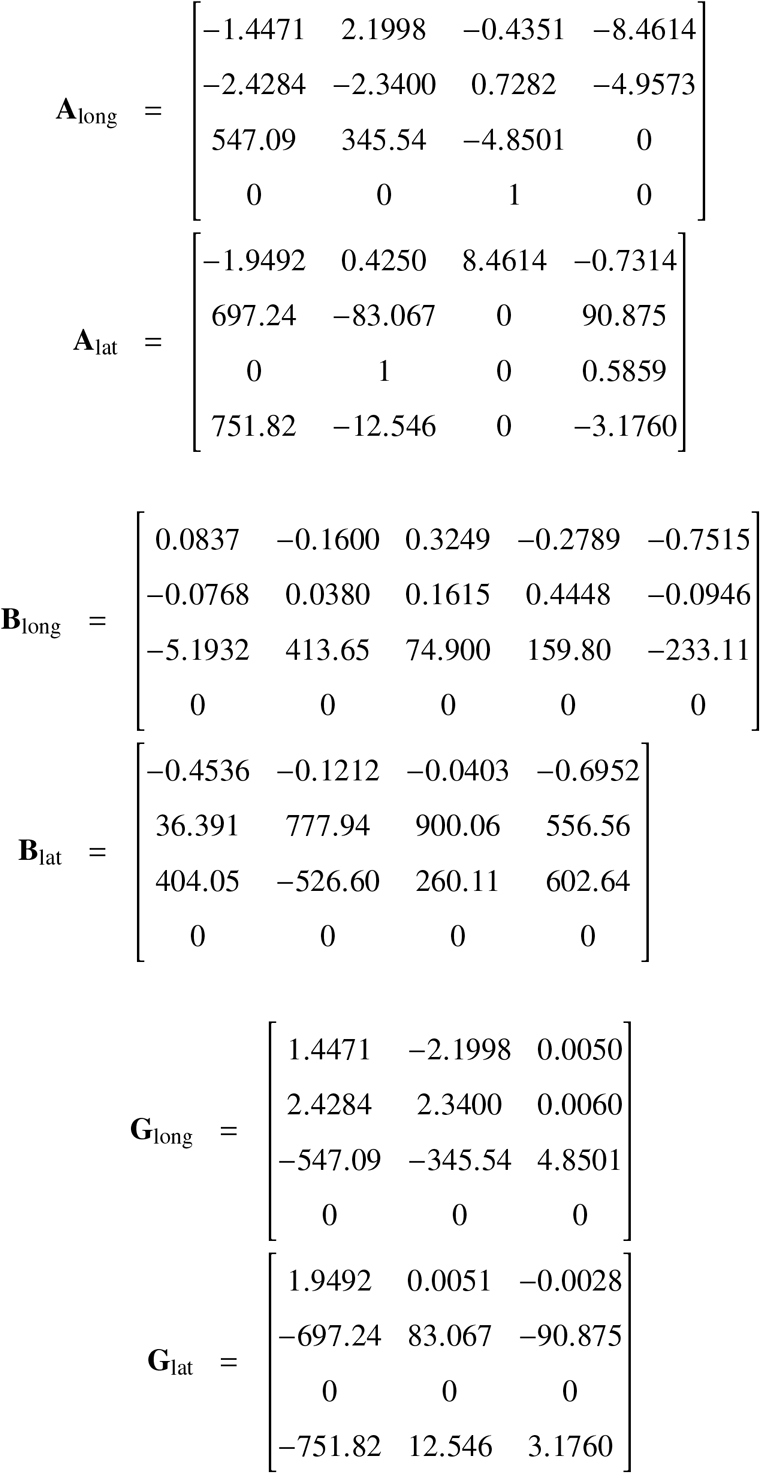

Writing the equations of motion in the body frame ℬ as above results in a simpler form than that which is obtained when using an axis system that is not necessarily aligned with the principal axes of the body. On the other hand, the visual axes of the head ℋ do not in general coincide with the principal axes of the body at equilibrium, and it is more reasonable to assume that the head is held level at equilibrium such that the retinal coordinates (0,0) coincide with the insect’s velocity vector in level flight at equilibrium. The set of body-fixed axes whose *x*-axis is aligned with the insect’s velocity vector at equilibrium, and whose transverse *y*-axis is normal to the insect’s plane of symmetry, is called the stability axis system 𝒮. In a final step, we therefore transform the equations of motion above from the body axes ℬ into the stability axes 𝒮, which simplifies the analysis in the main text. The details of these coordinate transformations are described further below.

### Coordinate Frames

Three different coordinate frames are employed in the preceding analyses (Fig. S2A): (i) the head (visual) axes 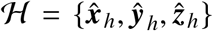, (ii) the body (principal) axes 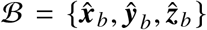, and the stability axes 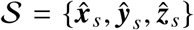. The visual ℋ axes and the body ℬ axes are defined anatomically, whereas the stability 𝒮 axes are specified by orienting the ***x***_*s*_ direction along the relative velocity vector for the given reference flight condition. The body axes ℬ are the natural coordinate frame in which to calculate the variations in the aerodynamic forces {*X, Y, Z* } and moments {*L, M, N*}, as the results can be straightforwardly transformed to stability axes 𝒮 for an arbitrary reference flight condition. Furthermore, because it assumed that the directions of the visual axes ℋ and the stability axes 𝒮 are aligned, the natural coordinate frame in which to express the flight dynamics and sensor outputs (Eqs. 1–2) is the stability frame 𝒮:

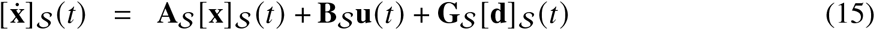

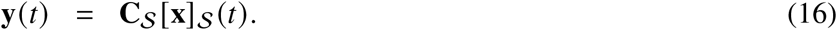

Bracket notation and subscripts are used to denote the coordinate frame for which a specific vector or matrix is expressed. In particular, [**x**]_*S*_ = [*δu*_*s*_ *δw*_*s*_ *δq*_*s*_ *δθ*_*s*_ *δv*_*s*_ *δ p*_*s*_ *δϕ*_*s*_ *δr*_*s*_]^*T*^ is the perturbation state expressed in stability coordinates, [**d**]_*S*_ = [*u*_*d,s*_ *w*_*d,s*_ *q*_*d,s*_ *v*_*d,s*_ *p*_*d,s*_ *r*_*d,s*_]^*T*^ are the gust inputs in stability coordinates, and the entries of the **A**_**𝒮**_, **B**_**𝒮**_, **G**_**𝒮**_, and **C**_**𝒮**_ system matrices are relative to the stability axes.

The electrophysiological characterization of LPTC response field properties was performed relative to the visual (head) axes ℋ, whose directions correspond to the stability axes 𝒮, therefore **C**_**𝒮**_ = **C**_**ℋ**_. The CFD characterizations of perturbation forces and moments (matrices **A**_**ℬ**_, **B**_**ℬ**_, and **G**_**ℬ**_) were generated relative to the body axes ℬ, so a transformation of the [**x**] _**ℬ**_ and [**d**] _**ℬ**_ vectors into stability axes 𝒮 is required to put the system into the form of Eqn. 15 to perform a comparison between the directions encoded by the LPTCs and the dynamically-significant directions defined by the Gramians.

To derive these transformations, we first consider the relationship between the velocity **v** and angular velocity **ω** in these two coordinate frames, which differ by a rotation *θ*_*e*_ about the pitch axis (Fig. S2):

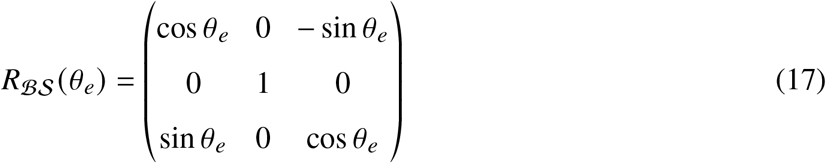

Hence, the velocity of the animal in body coordinates [**v**]_*B*_ is related to its velocity in stability coordinates [**v**]_*S*_ via

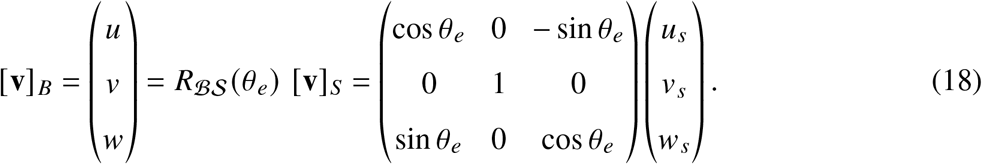

Similarly, the angular velocity of the animal in body coordinates [**ω**]_*B*_ is related to its velocity in stability coordinates [**ω**]_*S*_ via

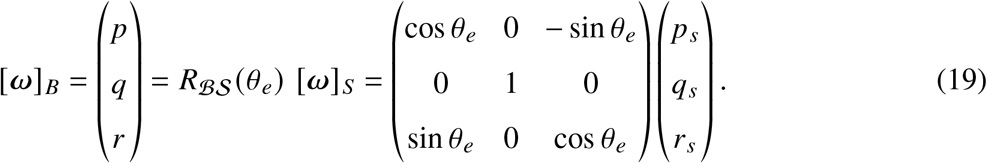

The full 8-dimensional state vector expressed in the body frame also contains two attitude angles, *θ* and *ϕ*. In the linearized flight dynamics these are actually perturbation angles from equilibrium, hence *θ* = *θ*_*s*_ since the 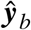 and 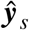 coordinate axes are colinear. To develop the relationship between *ϕ* and *ϕ*_*s*_, we take the axis-angle form for a rotation *ϕ*_*s*_ about the 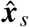 axis expressed in the 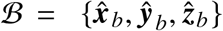 coordinate frame, convert this to its equivalent rotation matrix, then back out the equivalent 3-2-1 Euler angles {*ϕ, θ, ψ*} referred to the body ℬ frame. The axis-angle form is angle *ϕ*_*s*_ about axis 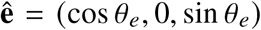. If we equate the corresponding rotation matrix for *ϕ*_*s*_ and 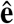 with the 3-2-1 Euler angle parameterization in body axes {*ϕ, θ, ψ*} of the same rotation, a ratio of the (3,2) and (3,3) entries results in

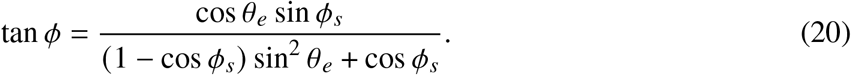

Hence for small *ϕ*_*s*_ and small *ϕ*, this reduces to *ϕ* ≈ (cos *θ*_*e*_)*ϕ*_*s*_.

Therefore, the longitudinal state transformation from stability 𝒮 to body axes ℬ is [**x**_long_]_ℬ_ = *R*_long_[**x**_long_]_𝒮_ is given by

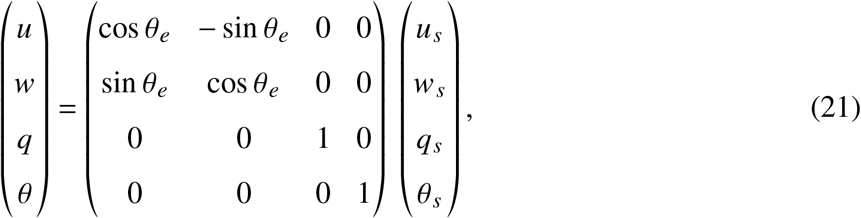

and the lateral state transformation from stability to body axes [**x**_lat_]_ℬ_ = *R*_lat_[**x**_lat_]_𝒮_ is given by

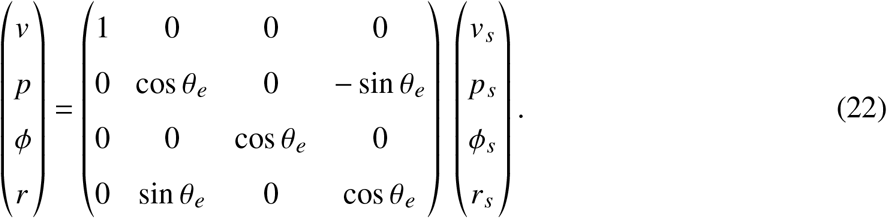

The transformation for the full state [**x**] _ℬ_ = *R*[**x**] _𝒮_ (longitudinal land lateral states combined) is

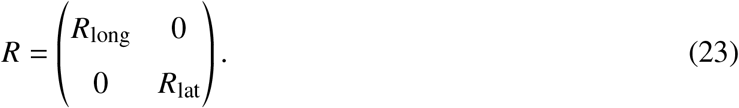

Since the disturbance vector **d** has six components compared to the eight components of the state vector **x**, we also define a transformation *R*_*G*_ which maps components of vectors expressed in the 𝒮 frame to the ℬ frame. This transformation is essentially the *R* matrix with the rows and columns associated with the *θ* and *ϕ* variables removed.

To express the **A**_ℬ_, **B**_ℬ_, and **G**_ℬ_ matrices relative to the stability axes 𝒮, we substitute 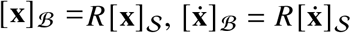 and [**d**] _ℬ_ = *R*_*G*_ [**d**] _𝒮_ into 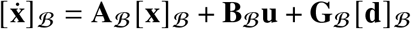,

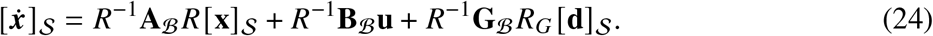

The final system matrices **A** = **A**_𝒮_, **B** = **B**_𝒮_, and **G** = **G**_𝒮_ (Eqn. 1), expressed in the stability axes 𝒮, are then **A**_𝒮_ = *R*^−1^**A**_ℬ_*R*, **B**_𝒮_ = *R*^−1^**B**_ℬ_, and **G**_𝒮_ = *R*^−1^**G**_ℬ_*R*_*G*_. To simplify our notation, we have omitted these S subscripts in the main text.

### Dynamically significant directions in state space

The Gramians constructed from the **A, B, G**, and **C** system matrices encode the dynamic properties of a system. These matrices have been widely used in testing for controllability and observability (*69–71*), in model reduction (*46, 70, 72, 73*), in sensor and actuator placement (*22, 74–76*), in disturbance rejection (*77, 78*), and in joint sensor and actuator design (*79, 80*). When the system matrix **A** is stable, the controllability and observability gramians are defined in the time domain as:

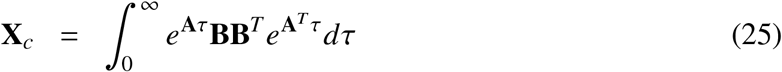

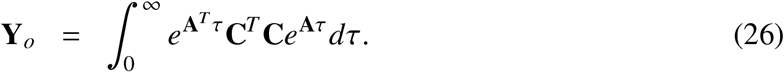

The disturbance sensitivity Gramian **X**_*d*_ is constructed similar to **X**_*c*_, by swapping the **B** with the **G** matrix in Eqn. 25. The Gramians are generated by computing the solutions to the Lyanunov equations,

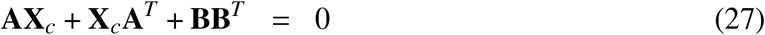

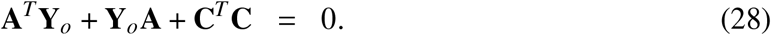

The controllability Gramian is related to the minimum signal energy required to reach a given state **x**_0_, which is given by 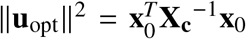. This is captured by the controllability ellipsoid 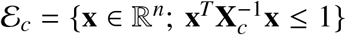 which defines the region of state space that can be reached by applying unit norm ∥**u**∥_2_ ≤ 1 input. The principal axes directions and lengths of ℰ_*c*_ are calculated as the eigenvectors and square roots of the eigenvalues of **X**_*c*_. Therefore, its larger axes represent the directions in state space requiring the lesser control effort to move along. Similarly, the disturbance sensitivity ellipsoid (ℰ_*d*_) is constructed in an identical fashion, but with **X**_*d*_ interchanged with **X**_*c*_. Its longest axes represent the directions of motion that are most readily excited by gusts. These most-sensitive directions characterise the worst-case disturbances that the insect may have to reject.

The observability Gramian is related to output energy for a given initial condition **x**_0_, since the energy of the output signal **y**(*t*) for an arbitrary **x**_0_ can be expressed as 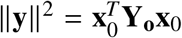. If we consider the set of initial conditions where ∥**y**∥^2^ ≤ 1, this generates the sensed directions in state space that have the smallest output norm. The observability ellipsoid 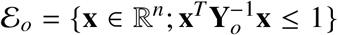 is created by replacing **Y**_*0*_ with its inverse 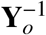, the covariance for the best unbiased estimate. Its principal axes correspond to directions in state space with the largest output norm, representing the specific self-motions that the insect is best-able to sense.

For system matrices **A** that are unstable, the integrals in Eqns. 25 and 26 are unbounded and **X**_*c*_ and **Y**_*0*_ are undefined. Generalizations to the controllability and observability Gramians were introduced in (*45*), corresponding to Eqns. 4 and 3 in the main text. For (**A, B**) stabilizable, (**C, A**) detectable, and no eigenvalues of **A** on the imaginary axis, the generalized Gramians are calculated as follows. First compute the solutions **P** and **Q** to the Riccati equations,

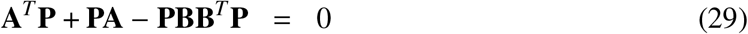

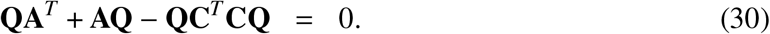

Next define **F** = −**B**^*T*^**P** and **L** = −**QC**^*T*^, and the generalized Gramians are the solutions to the Lyapunov equations,

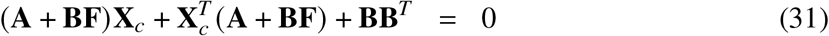

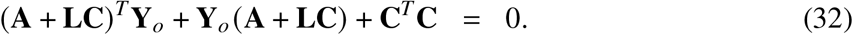

If the **A** matrix is stable, then **P** = 0 and **Q** = 0, and Eqns. 31 and 32 reduce to Eqns. 27 and 28. The authors in (*45*) also provide time domain interpretations of the generalized Gramians, and show they can be similarly related to the minimum control input energy 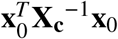, and average estimation error 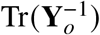.

Balanced realization theory for stable linear systems introduced by (*46*) provides additional tools for understanding the signal energy flow properties of a dynamical system. In particular, one can quantify the joint controllability and observability of a system that has been transformed into balanced coordinates 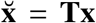, where 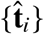 denote the columns of **T**^−1^. In these new coordinates the controllability and observability Gramians (Eqns. 25 and 26) are equal and diagonal, 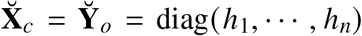, where the Hankel singular values 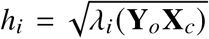 rank the joint controllability/observability of the direction 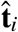 in original coordinates. In particular, directions with small *h*_*i*_ correspond to directions in state space that are simultaneously difficult to reach and observe, and contribute little to the overall behavior of the system. An extension to the unstable case was provided in (*45*), who showed that the same tools could be applied through the introduction of the generalized controllability Gramian **X**_*c*_ and observability Gramian **Y**_*0*_ as in Eqns. 4 and 3 in the main text. In particular, the Hankel singular values (HSVs) for an unstable system with no eigenvalues on the imaginary axis are the union of the HSVs for the partitioned stable and anti-stable subsystems, with the **A** matrix for the anti-stable partition replaced with −**A**, and these are computed similarly as 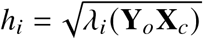. The transformation 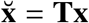 that was used to balance the system was based on the Cholesky factorization of **X**_*c*_ = **WW**^∗^. The eigenvalue decomposition of **W**^∗^**Y**_*0*_**W** = **VΣV**^∗^ is performed, with the resulting balancing transform computed as 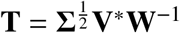.

### Supplementary results on the LPTC response fields

The V1-cell (Fig. 2C) shows a strong response to downward image motion over most of its frontolateral visual field, which rotates to become a preference for back-to-front image motion in the upper part of its caudolateral visual field (90° ≤ *γ* ≤ 150°), with vanishing sensitivity below the equator. This qualitative combination of response properties suggests that V1 will be most active during nose-up rotation. The V2-cell (Fig. 2D) responds strongly to upward image motion in the lateral visual field (45° ≤ *γ* ≤ 150°), which gradually changes into a preference for back-to-front image motion in the dorsofrontal visual field (0° ≤ *γ* ≤ 45°) and downwards image motion in the zone of binocular overlap (−30° ≤ *γ* ≤ 0°)(Fig. 2D). The right V2-cell will therefore be active during righthanded roll, with its preferred rotation axis directed ventral relative to the longitudinal roll axis (Fig. 2B). The Vx-cell (Fig. 2E) displays a preference for oblique upward image motion in the upper part of the frontal visual field (−30° ≤ *γ* ≤ 30°), which gradually changes into horizontal front-to-back motion (30° ≤ *γ* ≤ 105°) and becomes maximally sensitive to downward motion in the caudolateral visual field (105° ≤ *γ* ≤ 180°). Its preferred rotation axis is directed between the roll and pitch axes and tipped slightly ventrally, responding to rotations of the opposite sense to those that activate the ipsilateral V1- and V2-cells (Fig. 2G).

The response fields of the V1, V2 and Vx-cells in Fig. 2 plot only the ipsilateral parts of their response fields, including the narrow zone of binocular overlap. Some heterolateral LPTCs including V1 may respond to stimuli presented to the contralateral eye (*28, 31*), depending on the internal state of the fly (*32*). The same holds true for some of the VS-cells including VS6, which responds in a directionally-selective manner to stimuli presented to the contralateral eye (Fig. 2B). Though weaker than its ipsilateral motion sensitivity, this contralateral motion sensitivity enhances the cell’s response to roll motion. Inspection of the V1, V2 and Vx response fields (Fig. 2C-E) suggests that this is most likely to be attributable to excitatory input from the contralateral V2-cell. Along similar lines, simultaneous recordings have already established electrical connections between VS1-3 and the contralateral V1-cell (*81*), and ipsilateral inhibition of the VS1-cell by the Vx-cell (=Vi) (*28*). Simultaneous recordings have also demonstrated electrical coupling of neighbouring VS cells (*82*).

## Figures (SM)

**Figure S1:**
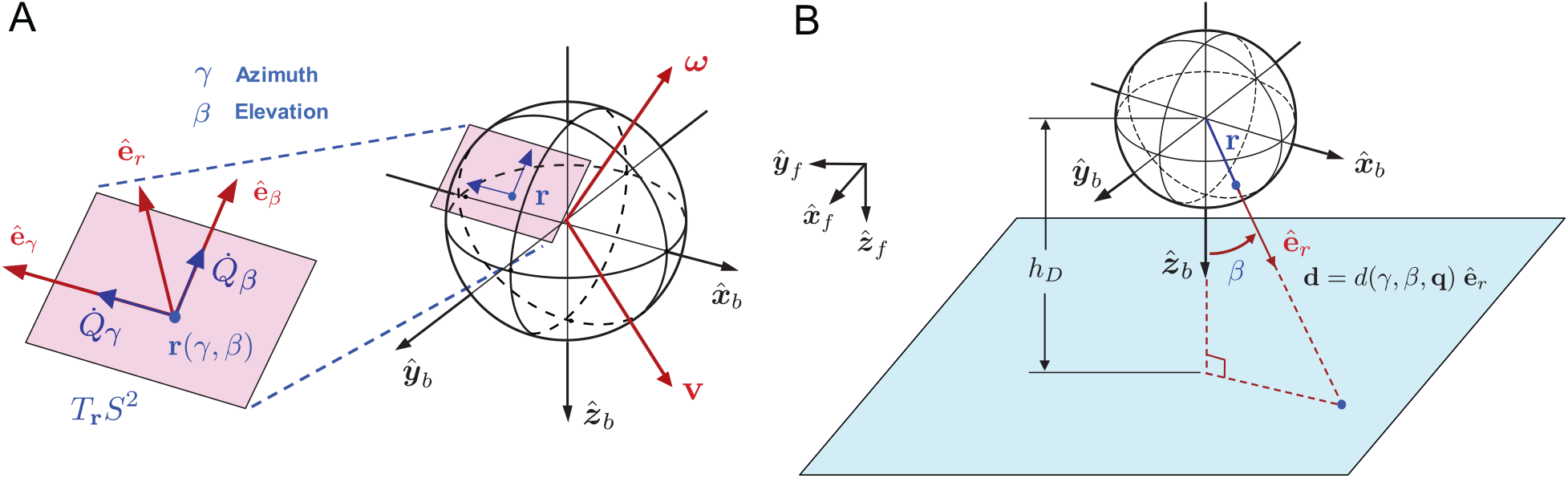
Geometry for spherical optic flow. (A) The azimuth 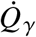 and elevation 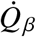 components of the optic flow vector are the projected relative velocities **ω** and **v** of visual contrasts or objects in the environment into the tangent space *T*_**r**_*S*^2^ of the imaging surface, modeled here as a sphere approximating the nearly 4*π* visual field of the compound eyes. (B) The function *d* (*γ, β*, **q**) is the distance from the imaging surface (**r**) to the nearest point in the environment.

**Figure S2:**
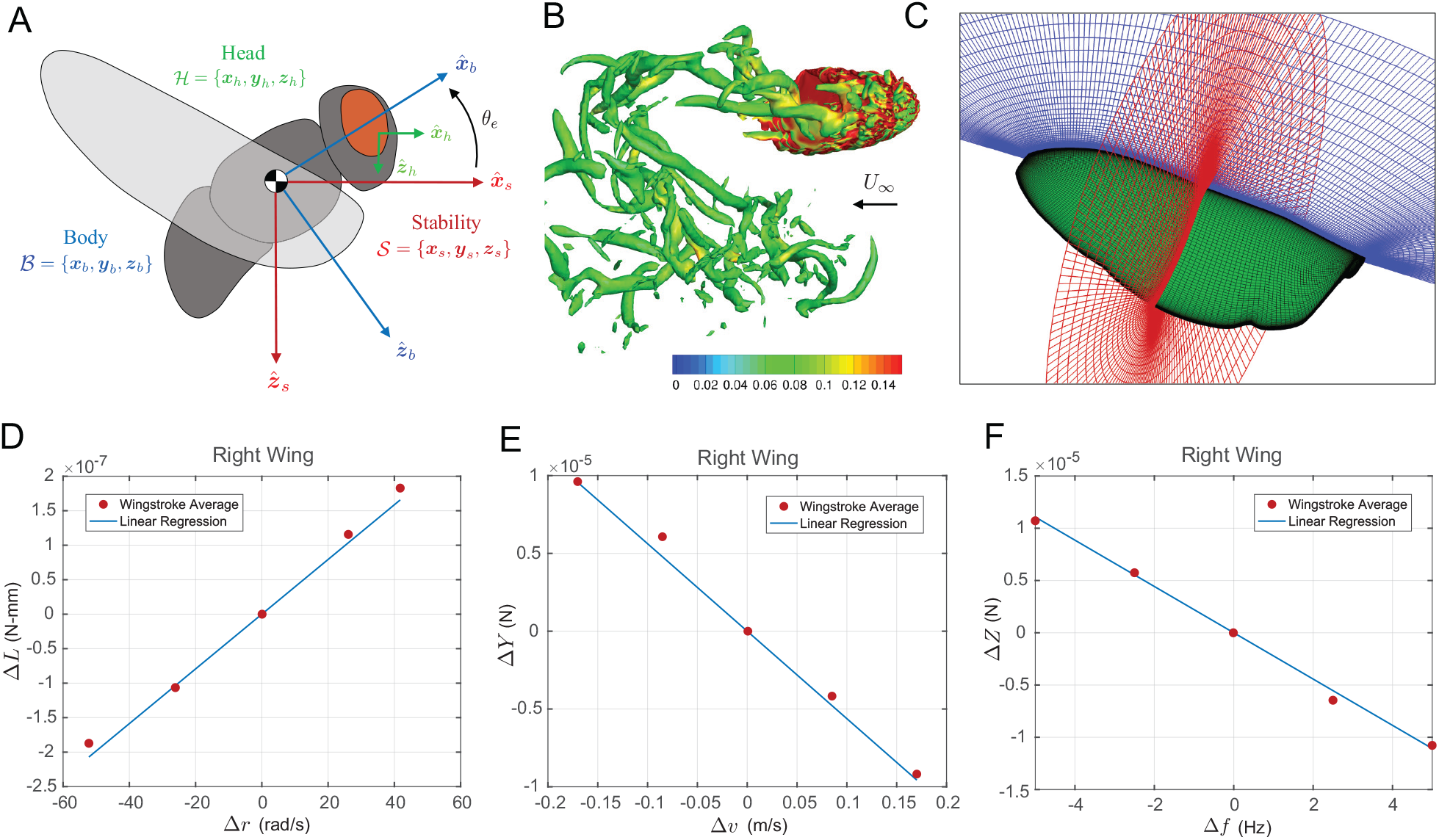
Reynolds Averaged Navier-Stokes (RANS) simulations for *Calliphora* wing motions. (A) Coordinate systems for analysis include the head (visual) axes 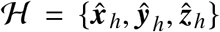, the body (principal) axes 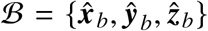, and the stability axes 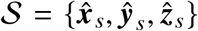. (B) Instantaneous wing wake structures are shown using iso-*Q* Criterion surfaces (*Q* = 0.001) and colored by vorticity at the end of the upstroke. (C) Simulations utilized an overset grid system for a body-fitted structured wing mesh and a Cartesian background mesh. (D-E) Example regressed wingstroke averaged aerodynamic moments and forces as a function of perturbation states and inputs for the right wing.

**Figure S3:**
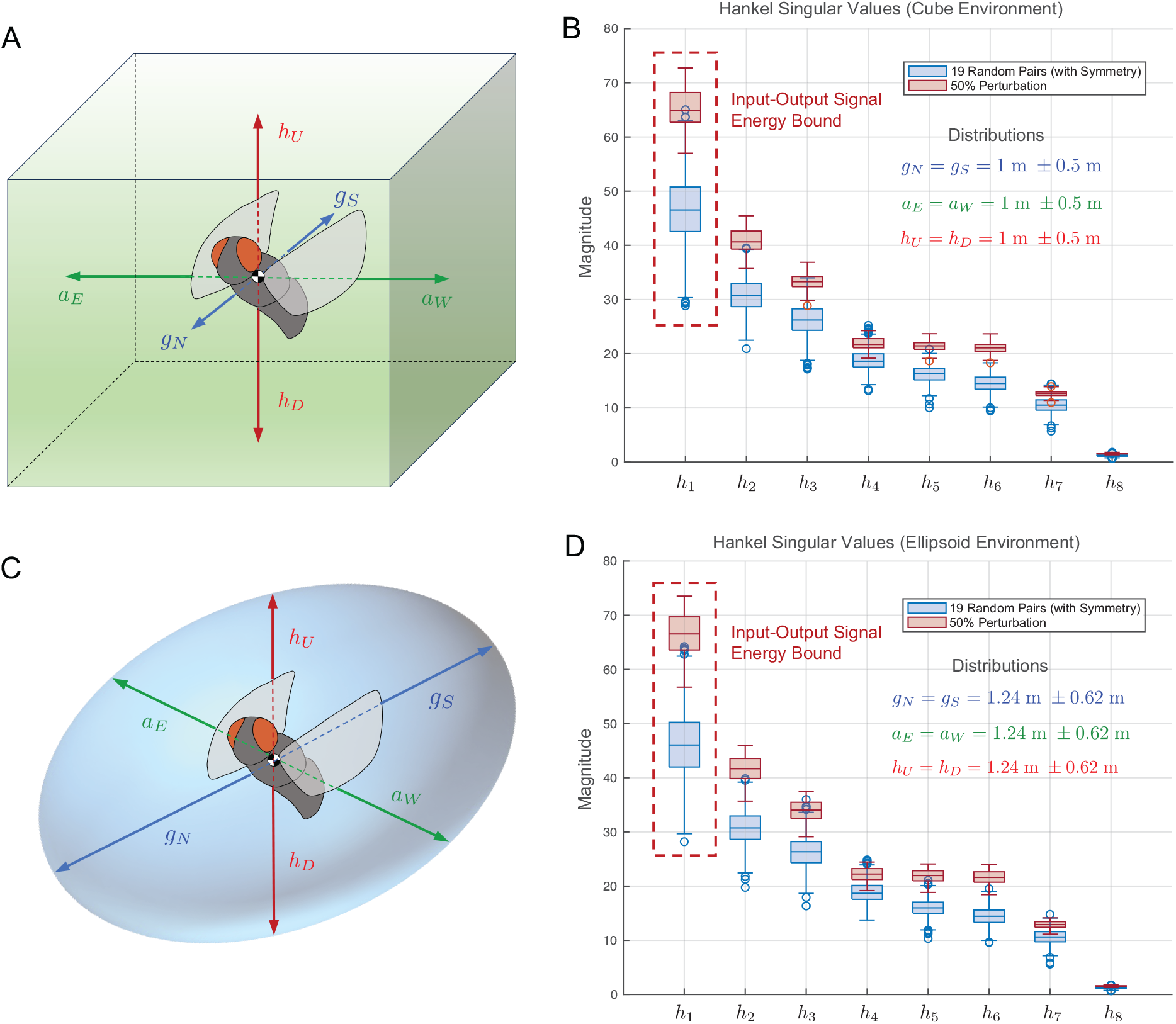
Robustness of Hankel singular values to environment perturbations. The dimensions (*g*_*N*_, *g*_*S*_, *a*_*E*_, *a*_*W*_, *h*_*u*_, *h*_*d*_) of two different nominal environment configurations with equal volume (cube, ellipsoid) are randomly selected from a uniform distribution (100 different configurations each) up to 50% and used to generate the system’s Hankel singular values. These distributions are compared to 1000 sets of randomly selected 19 left-right symmetric pairs from a uniform distribution on ℝ^8^. (A) Geometry of the cube environment. (B) Comparison of Hankel singular values for 100 randomly selected cube environment configurations. (C) Geometry of the ellipsoid environment. (D) Comparison of Hankel singular values for 100 randomly selected ellipsoid environment configurations.

**Figure S4:**
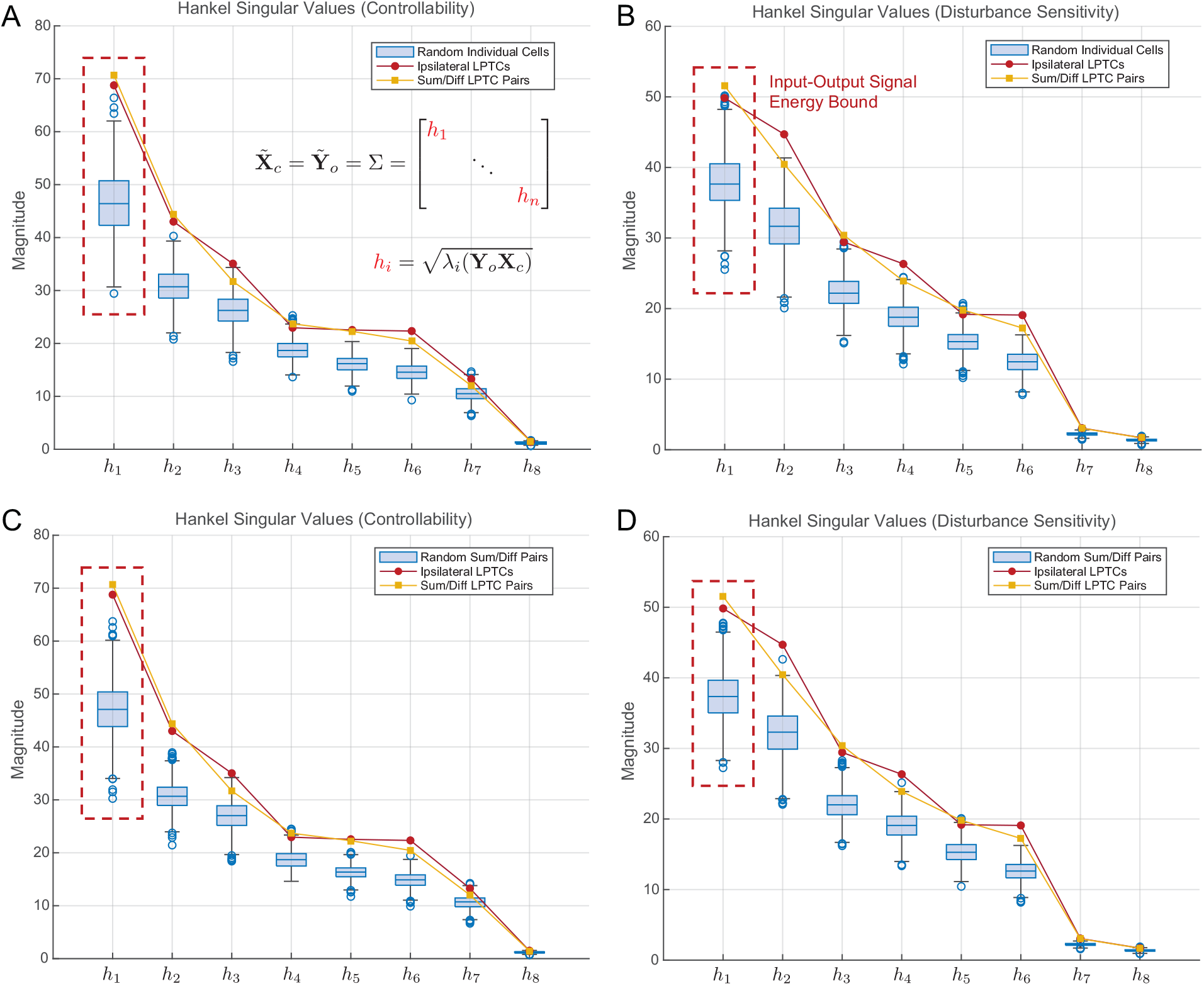
Robustness of Hankel singular values to environment perturbations. The dimensions (*g*_*N*_, *g*_*S*_, *a*_*E*_, *a*_*W*_, *h*_*u*_, *h*_*d*_) of two different nominal environment configurations with equal volume (cube, ellipsoid) are randomly selected from a uniform distribution (100 different configurations each) up to 50% and used to generate the system’s Hankel singular values. These distributions are compared to 1000 sets of randomly selected 19 left-right symmetric pairs from a uniform distribution on ℝ^8^. (A) Geometry of the cube environment. (B) Comparison of Hankel singular values for 100 randomly selected cube environment configurations. (C) Geometry of the ellipsoid environment. (D) Comparison of Hankel singular values for 100 randomly selected ellipsoid environment configurations.

**Figure S5:**
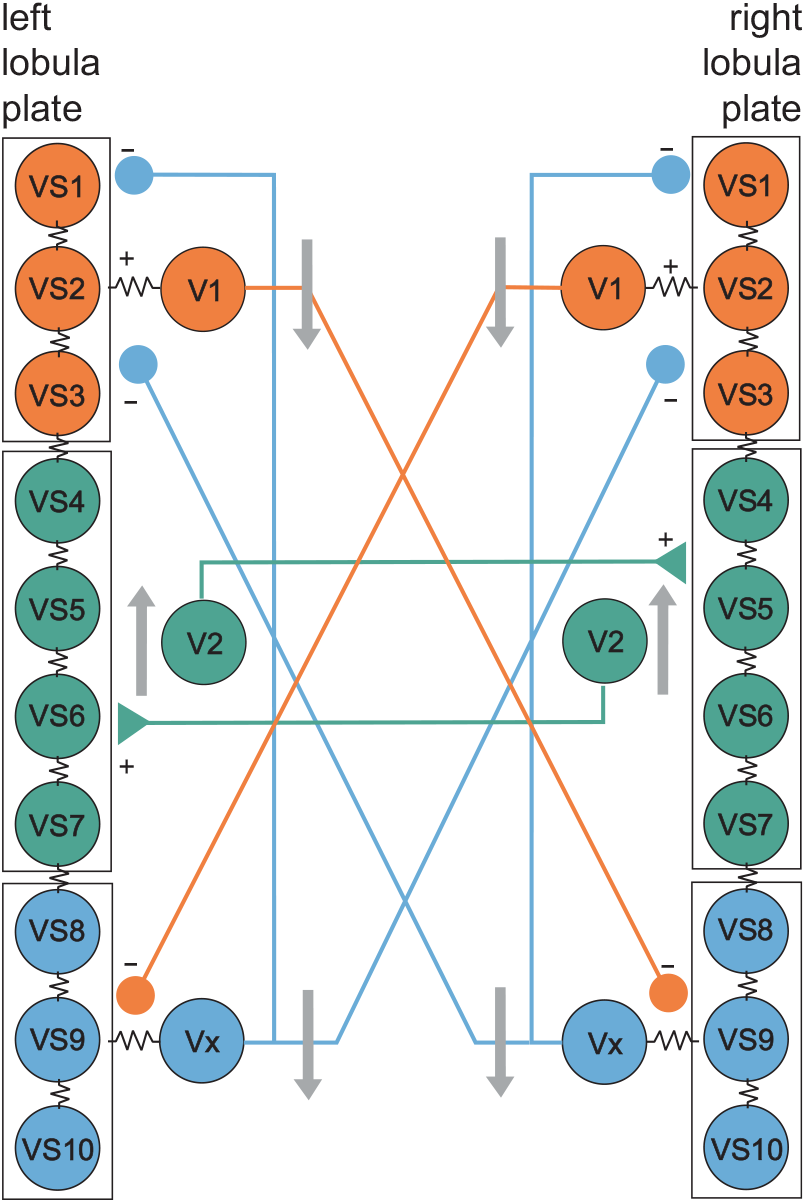
Putative synaptic coupling diagram of VS- and V-cells. Illustration of known (*83*) and hypothesized connections between the 10 VS-cells (VS1-VS10) and heterolateral V-cells (V1, V2, Vx) of the left and right lobula plate of the blowfly *Calliphora*. Colour coding corresponds to the proposed functional clusters shown in Fig. 2. Spherical and triangular symbols at the endpoints of connecting lines indicate inhibitory(−) and excitatory (+) outputs of the cells. Resistance symbols between cells refer to electrical synapses. Grey arrows indicate main sensitivity to vertical downward and upward motion of the LPTCs in the three clusters.

## Tables (SM)

**Table S1:** Moments of inertia about the principal axes of *Calliphora vicina* estimated from *μ*CT images.

| Insect | $m$ ($10^{-5}$ kg) | $I_1$ ($10^{-10}$ kg m<sup>2</sup>) | $I_2$ ($10^{-10}$ kg m<sup>2</sup>) | $I_3$ ($10^{-10}$ kg m<sup>2</sup>) |
| --- | --- | --- | --- | --- |
| male 1 | 6.792 | 5.802 | 5.730 | 1.457 |
| male 2 | 7.543 | 6.410 | 6.368 | 1.594 |
| female 1 | 7.953 | 6.924 | 6.883 | 1.822 |

**Table S2:** *Calliphora vicina* flapping wing simulation parameters.

|  |  |
| --- | --- |
| Freestream velocity $U_\infty$ (m s<sup>-1</sup>) | 0.8509 |
| Body pitch angle (°) $\theta_e$ | 30.365 |
| Nominal wing beat frequency (Hz) | 166.188 |
| Wing span (mm) | 10.1781 |
| Mean chord (mm) | 2.714 |
| Reynolds number | 1,746 |

**Table S3:** CFD Simulation setup parameters.

|  |  |
| --- | --- |
| Total timesteps | 2,880 (720 steps per cycle) |
| Flow condition | Laminar |
| Spatial reconstruction | 5<sup>th</sup> order WENO scheme |
| Temporal scheme | 2<sup>nd</sup> order backward difference (BDF2) |
| Low-Mach precondition | Yes |
| Overset grid node points | Wing (2.98 M) and Background (8.21 M) |

**Table S4:** Estimated stability derivatives (S.I. units).

| X Force ($10^{-5}$) | Z Force ($10^{-5}$) | M Moment ($10^{-7}$) |
| --- | --- | --- |
| $X_u = -8.3476$ | $Z_u = -14.009$ | $M_u = 2.2833$ |
| $X_w = 12.690$ | $Z_w = -13.499$ | $M_w = 1.4421$ |
| $X_q = -0.0286$ | $Z_q = -0.0345$ | $M_q = -0.0202$ |
| Y Force ($10^{-5}$) | L Moment ($10^{-7}$) | N Moment ($10^{-7}$) |
| $Y_v = -11.244$ | $L_v = 0.7428$ | $N_v = 3.1378$ |
| $Y_p = -0.0298$ | $L_p = -0.0884$ | $N_p = -0.0523$ |
| $Y_r = 0.0159$ | $L_r = 0.0968$ | $N_r = -0.0132$ |

**Table S5:**
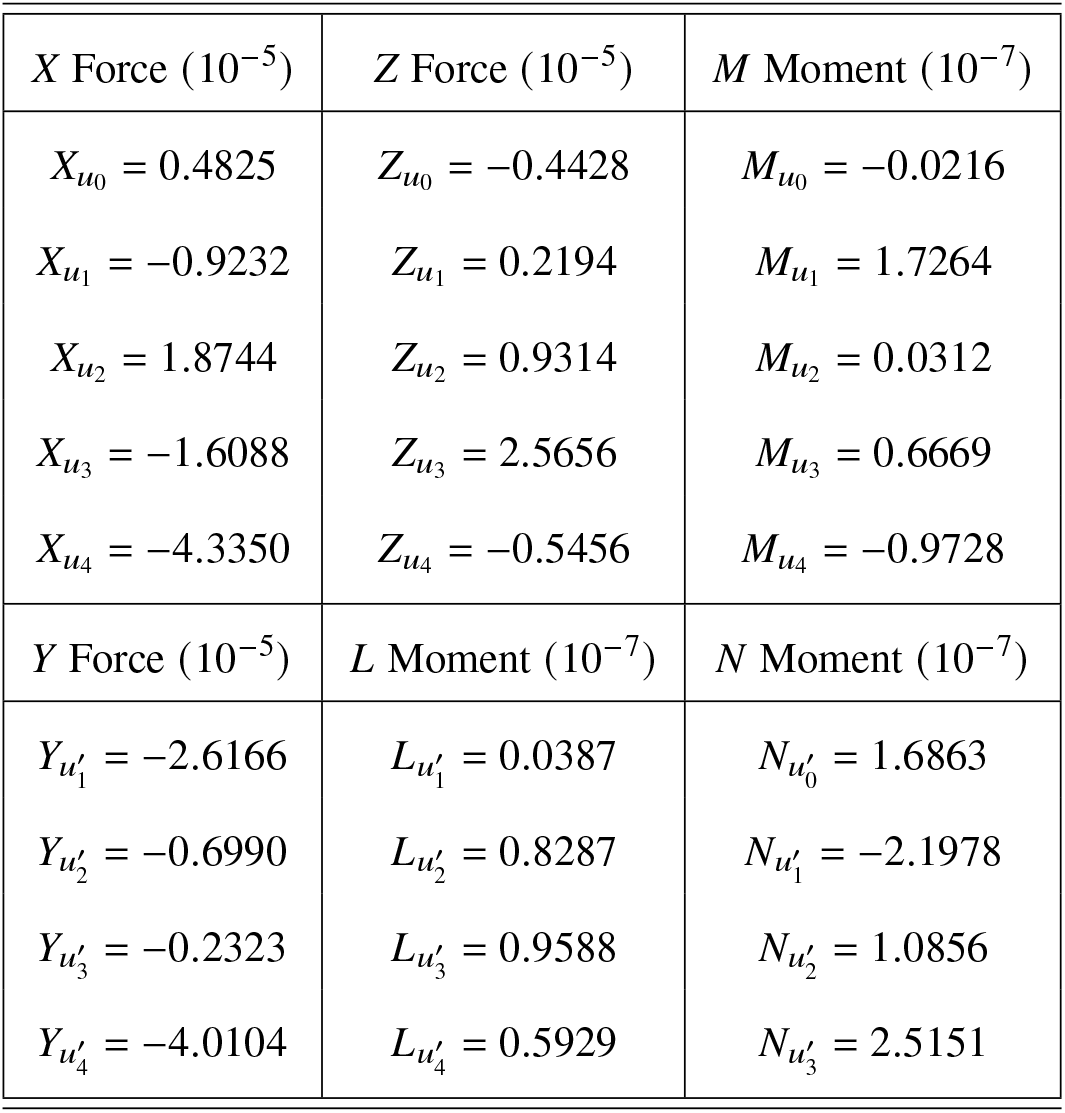
Estimated control derivatives (S.I. units).

| X Force ($10^{-5}$) | Z Force ($10^{-5}$) | M Moment ($10^{-7}$) |
| --- | --- | --- |
| $X_{u_0} = 0.4825$ | $Z_{u_0} = -0.4428$ | $M_{u_0} = -0.0216$ |
| $X_{u_1} = -0.9232$ | $Z_{u_1} = 0.2194$ | $M_{u_1} = 1.7264$ |
| $X_{u_2} = 1.8744$ | $Z_{u_2} = 0.9314$ | $M_{u_2} = 0.0312$ |
| $X_{u_3} = -1.6088$ | $Z_{u_3} = 2.5656$ | $M_{u_3} = 0.6669$ |
| $X_{u_4} = -4.3350$ | $Z_{u_4} = -0.5456$ | $M_{u_4} = -0.9728$ |
| Y Force ($10^{-5}$) | L Moment ($10^{-7}$) | N Moment ($10^{-7}$) |
| $Y_{u'_1} = -2.6166$ | $L_{u'_1} = 0.0387$ | $N_{u'_0} = 1.6863$ |
| $Y_{u'_2} = -0.6990$ | $L_{u'_2} = 0.8287$ | $N_{u'_1} = -2.1978$ |
| $Y_{u'_3} = -0.2323$ | $L_{u'_3} = 0.9588$ | $N_{u'_2} = 1.0856$ |
| $Y_{u'_4} = -4.0104$ | $L_{u'_4} = 0.5929$ | $N_{u'_3} = 2.5151$ |

**Table S6:** Eigenstructure of the symmetric and asymmetric modes of the system matrix **A** = **A**_S_, showing the eigenvalues (*λ*_*i*_) and eigenvectors for the symmetric and asymmetric parts, **x**_long_ = [*δu δw δq δθ*]^*T*^ and **x**_lat_ = [*δv δ p δϕ δr*]^*T*^, of the state vector **x** (S.I. units).

| symmetric | mode 1 | mode 2 | mode 3 | asymmetric | mode 4 | mode 5 | mode 6 |
| --- | --- | --- | --- | --- | --- | --- | --- |
| $\lambda_j$ | -19.89 | $6.77 \pm 14.92j$ | -2.28 | $\lambda_j$ | -63.38 | $-20.16 \pm 21.72$ | 15.51 |
| $\delta u$ | 0.0215 | $-0.0172 \pm 0.0244j$ | 0.0317 | $\delta v$ | -0.0017 | $0.0012 \pm 0.0149j$ | 0.0285 |
| $\delta w$ | 0.0506 | $-0.0200 \pm 0.0417j$ | -0.9042 | $\delta p$ | 0.9539 | $-0.9927 \pm 0.000j$ | 0.9852 |
| $\delta p$ | -0.9972 | $-0.9966 \pm 0.000j$ | 0.3900 | $\delta \phi$ | -0.0202 | $0.0306 \pm 0.0330j$ | 0.0853 |
| $\delta \theta$ | 0.0501 | $-0.0251 \pm 0.0554j$ | -0.1710 | $\delta r$ | -0.2993 | $-0.0891 \pm 0.0658j$ | 0.1458 |

**Table S7:**
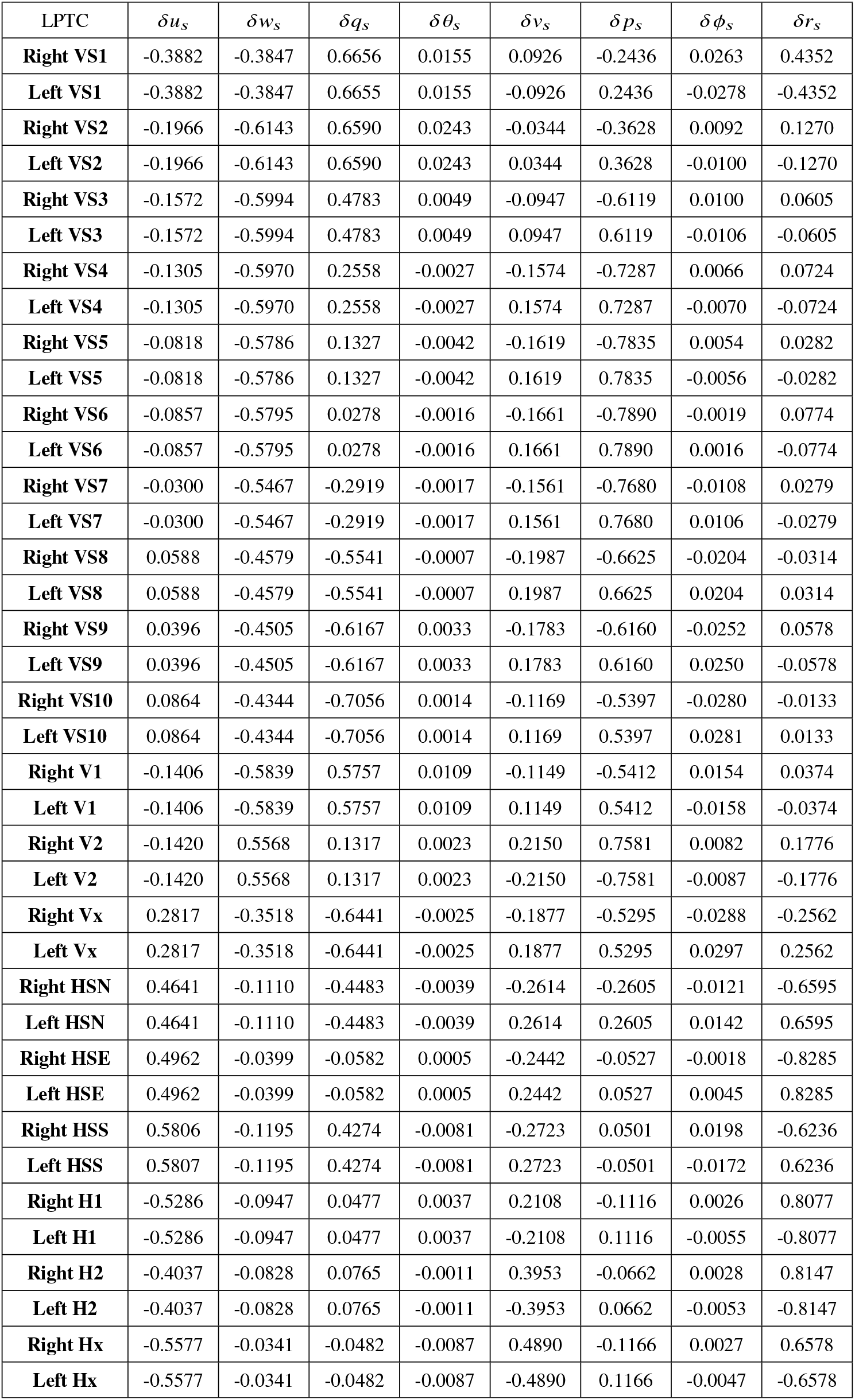
State encoding of the LPTC response fields expressed in the stability axes 𝒮 for the baseline enclosed rectangular prism. The rows of the unilateral output matrix **C**′ are formed from the elements corresponding to the motion state variables {*δu*_*s*_, *δw*_*s*_, *δq*_*s*_, *δθ*_*s*_, *δv*_*s*_, *δ p*_*s*_, *δϕ*_*s*_, *δr*_*s*_} scaled in S.I. units.

